# Chromatin-associated protein complexes link DNA base J and transcription termination in *Leishmania*

**DOI:** 10.1101/2020.05.26.117721

**Authors:** Bryan C Jensen, Isabelle Q. Phan, Jacquelyn R. McDonald, Aakash Sur, Mark A. Gillespie, Jeffrey A. Ranish, Marilyn Parsons, Peter J Myler

## Abstract

Unlike most other eukaryotes, *Leishmania* and other trypanosomatid protozoa have largely eschewed transcriptional control of gene expression; relying instead on post-transcriptional regulation of mRNAs derived from polycistronic transcription units (PTUs). In these parasites, a novel modified nucleotide base (β-D-glucopyranosyloxymethyluracil) known as J plays a critical role in ensuring that transcription termination occurs only at the end of each PTU, rather than at the polyadenylation sites of individual genes. To further understand the biology of J-associated processes, we used tandem affinity purification (TAP-tagging) and mass spectrometry to reveal proteins that interact with the glucosyltransferase performing the final step in J synthesis. These studies identified four proteins reminiscent of subunits in the PTW/PP1 complex that controls transcription termination in higher eukaryotes. Moreover, bioinformatic analyses identified the DNA-binding subunit of *Leishmania* PTW/PP1 as a novel J-binding protein (JBP3), which is also part of another complex containing proteins with domains suggestive of a role in chromatin modification/remodeling. Additionally, JBP3 associates (albeit transiently and/or indirectly) with the trypanosomatid equivalent of the PAF1 complex involved in regulation of transcription in other eukaryotes. Down-regulation of JBP3 expression levels in *Leishmania* resulted in a substantial increase in transcriptional read-through at the 3’ end of most PTUs. We propose that JBP3 recruits one or more of these complexes to the J-containing regions at the end of PTUs, where they halt progression of the RNA polymerase. This de-coupling of transcription termination from splicing of individual genes enables the parasites’ unique reliance on polycistronic transcription and post-transcriptional regulation of gene expression.

**Importance:** *Leishmania* parasites cause a variety of serious human diseases, with no effective vaccine and emerging resistance to current drug therapy. We have previously shown that a novel DNA base called J is critical for transcription termination at the ends of the polycistronic gene clusters that are a hallmark of *Leishmania* and related trypanosomatids. Here, we describe a new J-binding protein (JBP3) associated with three different protein complexes that are reminiscent to those involved in control of transcription in other eukaryotes. However, the parasite complexes have been reprogrammed to regulate transcription and gene expression in trypanosomatids differently than in the mammalian hosts, providing new opportunities to develop novel chemotherapeutic treatments against these important pathogens.

## Introduction

The genus *Leishmania* includes several species of protozoan parasites that cause a spectrum of human diseases, ranging from cutaneous lesions to disfiguring mucocutaneous and lethal visceral leishmaniasis, depending primarily on the species involved. *Leishmania* is transmitted through the bite of the sand fly and belongs to the family Trypanosomatidae, which also includes the vector-borne human pathogens *Trypanosoma brucei spp*., causative agent of human African trypanosomiasis (African sleeping sickness), and *Trypanosoma cruzi*, causative agent of Chagas’ disease. Reflecting the ancient divergence of these organisms, the Trypanosomatidae exhibit a myriad of biological differences from “higher” eukaryotes. One major difference is that each chromosome is organized into a small number of polycistronic transcription units (PTUs), which consist of tens-to-hundreds of protein-coding genes co-transcribed from a single initiation site at the 5’ end of the PTU to a termination site at the 3’ end (1). Interestingly, unlike the operons of prokaryotes, genes within each PTU are not confined to a single pathway or function. Individual genes within the primary transcript are *trans*-spliced by addition of a 39-nucleotide spliced-leader (SL) mini-exon to provide the 5’ cap-4 structure and polyadenylated to form the mature individual mRNAs. As a result of this unique genomic organization, all genes within a PTU are transcribed at the same rate (2, 3). Hence, gene expression must be controlled by post transcriptional processes, such as splicing/polyadenylation rate, RNA stability and translational regulation.

A second distinct feature of the Trypanosomatidae (and other Euglenazoa) is that ∼1% of the thymidine bases in the nuclear genome are glucosylated to form the novel nucleotide β-D-glucopyranosyloxymethyluracil, usually referred to as J (4, 5). While the majority of J is localized within telomeric repeat sequences (4, 6-9), chromosome-internal J is found at almost all transcription termination sites (10, 11) and centromeres, which also correspond to the major replication origins on *Leishmania* chromosomes (12, 13). In some trypanosomatids (although not *Leishmania*), J is also found in transcriptionally silent regions containing retrotransposons and/or other repetitive sequences (9). J biosynthesis occurs in two steps, whereby one of two proteins (JBP1 or JBP2) hydroxylates the methyl group of thymidine to form hydroxymethyldeoxyuracil (HmdU), which is subsequently further modified by a glucosyltransferase (HmdUGT) to form J (10, 14). Both JBP1 and JBP2 have an N-terminal oxygenase (Tet_JBP) catalytic domain; but JBP1 contains a central J-binding domain (15), while JBP2 instead contains SNF2_N, ATP-binding and helicase C-terminal domains that are suggestive of a role in chromatin binding and/or re-modeling (16, 17). Null mutants of *jbp2* have not been isolated (despite multiple attempts) in *Leishmania*, suggesting that it is an essential gene (and hence that J is required for viability) (18). In contrast, *jbp2* null mutants have been isolated and shown to have 40% less J than wild-type parasites (11). Chromosome-internal J was gradually lost during continuous growth of *Leishmania tarentolae jbp2* null mutants, with a concomitant increase in read-through transcription at termination sites, suggesting a critical role for J (and JBP2) in transcription termination. Coupled with the identification of a JBP1 recognition motif (19), these data led to the model that JBP1 is responsible for J maintenance after DNA replication by binding to pre-existing J on the parental strand and modifying the thymidine twelve nucleotides downstream on the newly synthesized strand. Conversely, JBP2 is proposed to be largely responsible for *de novo* synthesis of J only when JBP1 is not able to fully restore J on both strands.

Despite this knowledge of the enzymes involved in J biosynthesis, we currently know little about how J mediates transcription termination (and/or repression of initiation), since the machinery underlying this process has not yet been identified. Here, we have used Tandem Affinity Purification (TAP)-tagging and tandem mass-spectrometry to identify a PTW/PP1-like protein complex that interacts with HmdUGT. This complex includes a novel J-binding protein (JBP3) that appears to be essential in *Leishmania*. RNA-seq analysis following conditional down-regulation of JBP3 expression shows substantially higher levels of transcriptional read-through at the 3’ end of most PTUs, suggesting that it plays an important role in transcription termination. While this manuscript was being prepared, Kieft et al (20) reported similar results in *Leishmania* and *T. brucei*. Here, we have extended these findings by demonstrating that JBP3 also interacts with another protein complex likely involved in chromatin modification/remodeling, as well as to a lesser degree with a PAF1C-like complex that likely interacts with RNA polymerase II (RNAP II). Therefore, despite the differences in gene regulation from other eukaryotes, *Leishmania* appears to utilize proteins related to those used for chromatin remodeling and transcriptional regulation in other eukaryotes to provide the molecular machinery that links J to termination of RNAP II-mediated transcription.

## Results

### Identification of a protein complex containing a novel J-binding protein

To date only three proteins have been shown to be involved in J biosynthesis: JBP1, JBP2 and HmdUGT (hereafter referred to as GT). To expand the network of proteins important in J biosynthesis and/or function, we used mass spectrometry to identify proteins that co-purified with a TAP-tagged GT bait expressed in *L. tarentolae*. Two separate experiments were performed: the first used extracts from wild-type parasites constitutively expressing the tagged protein, while the second pull-down (and all other TAP-tag experiments) were performed using extracts from a tetracycline (Tet) induced T7-TR cell line, which over-expresses the tagged protein integrated at the *ODC* locus (see **Fig. S1A** in the supplemental material). After affinity purification, SDS-PAGE and silver staining confirmed successful enrichment of the bait protein in the pooled eluates (**Fig. 1A)**. Proteins were identified by liquid chromatography-tandem mass spectrometry (LC-MS/MS) and their (log_2_-fold) enrichment calculated by comparison to a control (untransfected) parental cell line (see **Data Set S1** in the supplemental material for a complete list of all proteins detected in each TAP-tag pull-down).

**FIG 1.**
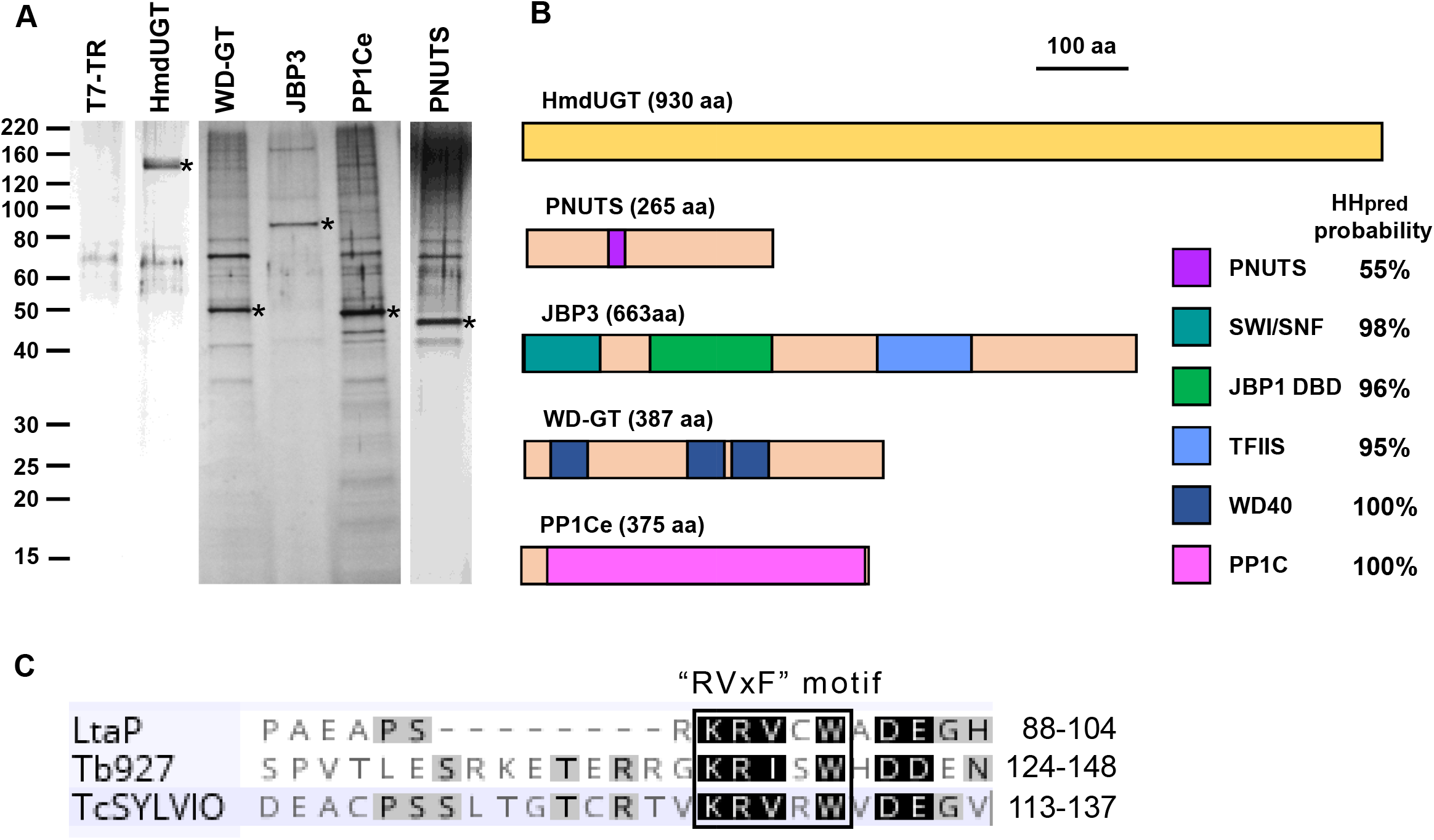
The PJW/PP1 complex. **A**. Proteins in the peak fractions from TAP purifications of HmdUGT and other components of the PJW/PP1 complex (WD-GT, JBP3, PP1Ce and PNUTS) overexpressed in T7-TR cells were separated by 4-20% SDS-PAGE and silver stained. Each lane represents 5% of the total fraction. The TAP-tagged “bait” protein is indicated by an asterisk. The first lane shows the equivalent fraction from a mock purification of control (T7-TR) cells. **B**. Schematic representation of key domains in the four proteins (and GT) in the PJW/PP1 complex, as predicted by HHpred analysis. The probability of the match between the *Leishmania* protein and the most similar experimentally determined structure is shown to the right. Potential functions of each domain are discussed in the text. **C**. The putative PP1C-interacting domain of *L. donovani* (LtaP), *T. brucei* (Tb927) and *T. cruzi* (TcSYLVIO) PNUTS is shown and the “RvxF” docking motif (which has consensus sequence of K/R-K/R-V/I-X-F/W, where X is residue other than F, I, M, Y, D or P)*(21)*, is shown above. The amino acid postions for each end of the domain are shown to the right.

The data revealed substantial (>500-fold) enrichment of five proteins (including the “bait”) in the GT-TAP pulldown; four that were enriched in both replicates and one that was detected only in the first experiment (**Table 1)**. These results suggested that the five proteins form a GT-associated protein complex, which was investigated in more detail, as described below. In addition, six prefoldin subunits were enriched by 50-to 450-fold in one or both replicates (see **Table S1** in the supplemental material, which includes all proteins that were substantially enriched in each pulldown). However, since prefoldin likely acts only as a chaperone for one or more proteins of the GT-associated complex, these proteins will not be further considered here.

**Table 1.** Enrichment of proteins in the PJW/PP1 complex.

| Gene ID | Name | HHpred/InterPro domains | Tagged protein |  |  |  |  |
| --- | --- | --- | --- | --- | --- | --- | --- |
|  |  |  | HmdUGT | PP1Ce | PNUTS | JBP3 | WD-GT |
| LtaP36.2450 | HmdUGT | None found | 14.1 | 7.2 | 8.2 | 10.6 | 9.2 |
| LtaP15.0230 | PP1Ce | Serine/threonine-protein phosphatase | 7.4 | 17.1 | 12.2 | 10.9 | 13.9 |
| LtaP33.1440 | PNUTS | PPP1R10/PNUTS | (9.8) | 12.5 | 17.7 | 10.3 | 13.2 |
| LtaP36.0380 | JBP3 | SWI1, J-binding & N-terminal TFIIIS | 11.1 | 12.3 | 10.3 | 15.9 | 12.5 |
| LtaP32.3990 | WD-GT | WD40 repeats | 9.3 | 14.8 | 12.7 | 12.3 | 16.3 |

Enrichment is expressed as the mean log<sub>2</sub> fold-change compared to control pull-downs.
Parentheses indicate that enrichment was observed in only one replicate.

One of the highly enriched proteins in the GT-TAP pulldown (LtaP15.0230) was annotated in TriTrypDB as a putative protein phosphatase 1 catalytic subunit, while the other three (LtaP36.0380, LtaP33.1440 and LtaP32.3990) were all annotated as “Hypothetical protein, conserved”. Constructs were made for expression of TAP-tagged proteins and transfected into the *L. tarentolae* T7-TR strain. Affinity purification was performed on two (independently generated) cell lines for each version of the TAP-tagged proteins (the results from one replicate of each are shown in **Fig. S1B**) and analyzed by LC-MS/MS (see **Data Set 1**). The results from these pull-downs (**Table 1** and **Table S1**) show the four proteins identified above were all highly enriched (as was GT, albeit to a lesser degree). Thus, we conclude that these proteins form a stable complex, with GT perhaps more transiently associated than the other four components.

*LtaP15*.*0230* encodes one of eight isoforms of the catalytic subunit of protein phosphatase 1 (PP1C) found in the *L. tarentolae* genome. Phylogenetic analysis (see **Fig. S2**) indicates that there are five different clades of PP1C paralogues in trypanosomatids and PP1Ce (encoded by *LtaP15*.*0230*) belongs to a clade that lacks any mammalian orthologue. Interestingly, Salivarian trypanosomes (including *T. brucei*) also lack PP1Ce, although it is present in Stercorarian trypanosomes (**Fig. S2**) and the more distantly related *Blechomonas ayalai, Paratrypanosoma confusum* and *Bodo saltans* (data not shown). TAP-tagging of PP1Ce resulted in co-purification of seven proteins with >150-fold enrichment (**Table S1**). These include GT and the other three components of the complex described above, as well as three proteins (encoded by *LtaP05*.*1290, LtaP07*.*0770* and *LtaP29*.*0170*) annotated as protein phosphatase regulatory subunits (PPP1R7/Sds22, PPP1R11/inhibitor 3, and PPP1R2/inhibitor 2, respectively). In higher eukaryotes PPP1R2 and PPP1R11 use the same “RVxF” docking motif to bind to and inhibit PP1C, while PPPR7 docks at a different site *(21)*. At least some mammalian PP1C proteins form an inactive heterotrimeric complex containing PP1R7 and PPP1R11 (*22*). Thus, we suggest that PP1Ce forms at least three separate complexes; one with both PPP1R7 and PPP1R11, a second with PPP1R2 (which may or may not contain PPP1R7), and the third being the GT-associated complex, described below.

We performed a series of bioinformatic analyses to identify domains and/or motifs that might provide hints as to the function of the three proteins of GT-associated complex that lack a functional description. BlastP and InterProScan searches showed high confidence matches only to orthologues in other trypanosomatids with no informative domains identified. However, HHpred analysis, which detects remote protein homology by using hidden Markov models and structure prediction (23), revealed a number of structural matches, that are summarized by the cartoons in **Fig. 1B** and shown in detail in **Fig. S3**. HHpred predicts that the central portion (residues 88-105) of LtaP33.1440 contains structural similarity to a portion of the human serine/threonine-protein phosphatase 1 regulatory subunit 10 (PPP1R10), also known as PNUTS (for PP1 nuclear targeting subunit) (24-27). The trypanosomatid protein is much smaller (264 *vs* 940 amino acids) than mammalian PNUTS, with the sequence and structural similarity restricted to the central region above (**Fig. S3A**). However, this region contains the “RVxF” sequence motif (**Fig. 1C**) noted above, which is present in most inhibitors and responsible for interaction with PP1C (28). Therefore, in deference to precedent in the field (20), we will refer to this protein as PNUTS, despite the lack of an obvious nuclear localization signal. TAP-tagging of PNUTS showed significant enrichment of the four other components of the GT-associated protein complex and no other proteins (**Table S1**).

BlastP searches of LtaP32.3990 returned matches to orthologues in other trypanosomatids, as well as WD40 repeats in proteins from several other organisms, while HHpred analysis (**Fig. S3B**) identified at least three WD40 repeats. Therefore, we will refer to this protein as WD-GT, to distinguish it from the numerous other WD40 repeat-containing proteins in *Leishmania*. TAP-tagged WD-GT pulled down the four other components of the GT-associated protein complex; as well as a number of chaperone-associated proteins, including prefoldins, T-complex and heat shock proteins (**Table S1**).

HHpred analysis of LtaP36.0380 revealed three separate domains with structural similarity to different proteins (**Fig. S3C**). The N-terminal domain (residues 2-86) is similar to the central portion of the SWI1 subunit of the yeast SWI/SNF chromatin remodeling complex, while the C-terminal domain (residues 384-485) is related to the N-terminal TFIIS domain of mammalian PNUTS. Most importantly, the central portion (residues 137 to 269) is predicted to have substantial structural similarity to the DNA-binding domain (DBD) of JBP1 and *in silico* folding of this region revealed conservation of the functional signature D-(W/F/Y)-x-x-GGTRY motif present in all trypanosomatid JBP1 proteins (**Fig. 2A**). In addition, a structural model of the DBD from LtaP36.0380 contains a binding pocket large enough to accommodate the glucose ring of J (**Fig. 2B and C**) and preliminary experiments indicate that it binds preferentially to J (personal communication from Anastasis Perrakis at NKI, Amsterdam). While this manuscript was in preparation, this J-binding function of LtaP36.0380 was experimentally confirmed by others (20), so the protein was re-named <u>J</u>-<u>b</u>inding <u>p</u>rotein 3 (JBP3).

**FIG 2.**
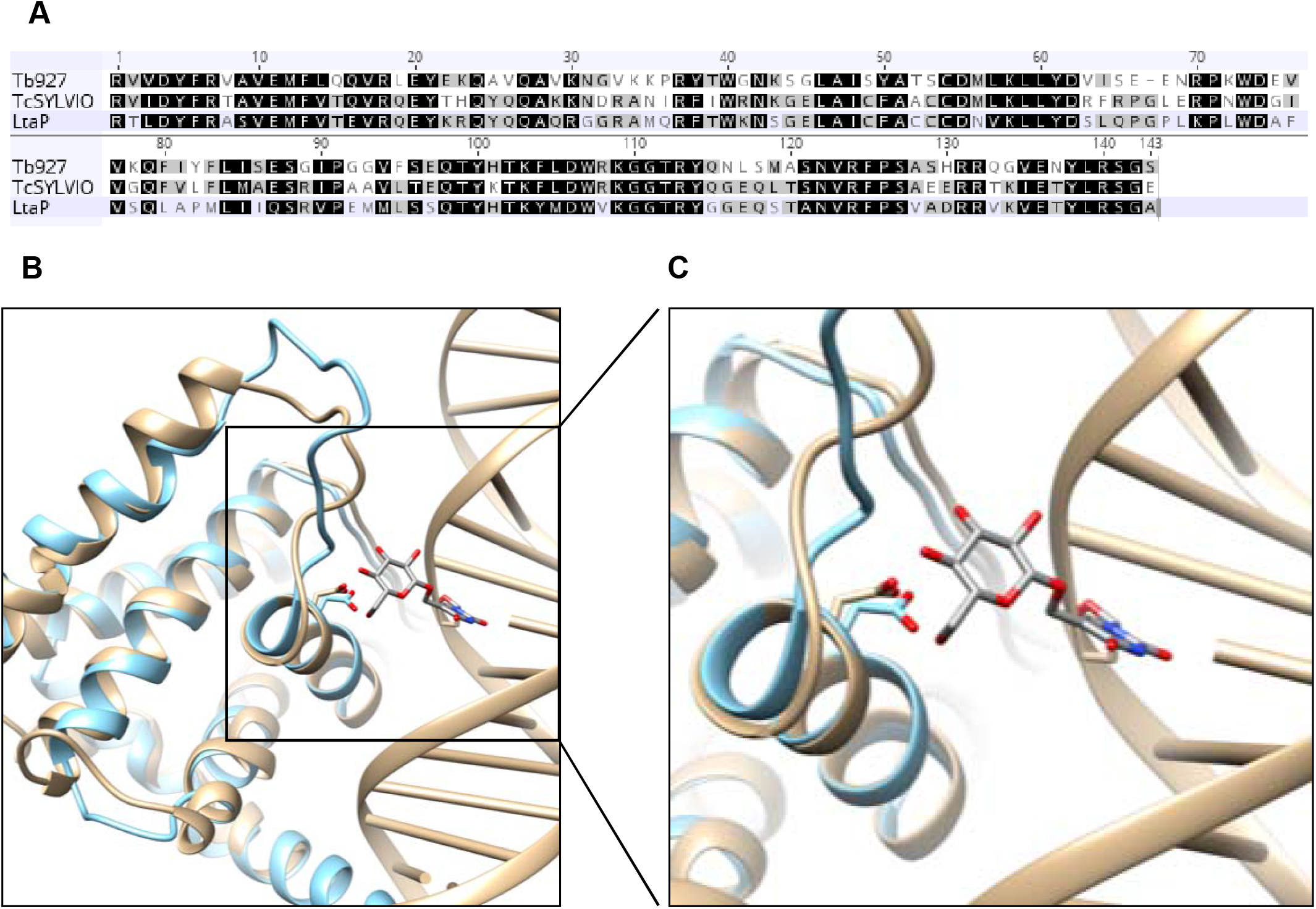
Modeling of the JBP3 DNA-binding domain. **A**. Sequence alignment of the putative JBP3 J-binding domain from *T. brucei* EATRO927 strain, *T. cruzi* Silvio strain, and *L. tarentolae* Parrot strain. Residues that are identical or conservatively replaced in all three species are shaded black, while those that are identical or conserved in two species are shaded grey. **B**. The structure of the DNA binding domain from JBP3 (light blue) was modeled using RosettaCM against the J-binding domain of JBP1 (tan) from PDB entry 2XSE. The interaction between the conserved aspartic acid residue (Asp_525_ in JBP1 and Asp_241_ in JBP3-) with the glucose of base J is shown. **C**. Higher resolution view of the of the interaction between conserved aspartate of both proteins and base J.

The molecular characteristics of the four proteins identified in the GT pull-downs – a PP1 catalytic subunit (PP1Ce), a predicted PP1 regulatory protein (PNUTS), a WD40 repeat protein (WD-GT), and a DNA binding protein (JBP3) – are highly reminiscent of the components of the mammalian PTW/PP1 complex. This complex, which contains PP1C, PNUTS, WDR82, and the DNA-binding protein TOX4, has a role in controlling chromatin structure (27, 29). Importantly, the mammalian PTW/PP1 complex was recently found to negatively regulate RNAP II elongation rate by dephosphorylating transcription elongation factor Spt5, leading to transcription termination at polyadenylation sites (30). Thus, our results indicate that GT associates with a PTW/PP1-like complex in *Leishmania* (which we will refer to as PJW/PP1), wherein JBP3 replaces the DNA-binding function of TOX4.

### JBP3 is part of another chromatin remodeling complex

While tandem affinity purification of TAP-tagged JBP3 showed >256-fold enrichment of the PJW/PP1 complex proteins (PP1Ce, PNUTS, WD-GT, and GT)(**Table S1**), another four proteins (encoded by *LtaP35*.*2400, LtaP28*.*2640, LtaP12*.*0900* and *LtaP14*.*0150*) were >3000-fold enriched (**Table 2**). BlastP analyses of these proteins failed to reveal convincing matches to anything other than orthologues in other trypanosomatids and InterProScan showed no matches with the default parameters. However, LtaP35.2400 is annotated as a “SET domain-containing protein, putative” in TriTrypDB and HHpred analysis revealed that the N-terminal region (amino acids 70-227) contains structural similarity to SET domain-containing proteins, while the central portion (residues 355-385) shows weaker similarity to C4-type Zinc finger domains from several unrelated proteins (**Fig. S3D**). SET domains, which are usually involved in binding to and/or methylation of histones (31), are also present in several other *Leishmania* proteins, so we have named this protein SET-J3C to distinguish it from the others. HHpred analysis of LtaP14.0150 showed structural similarity to Chromatin organization modifier (Chromo) domains (*32, 33*) from numerous eukaryotic proteins at its N-terminus (amino acids 1-55) and weak similarity to Chromo shadow domains at the C-terminus (34) (**Fig. S3E**).

**Table 2.** Enrichment of proteins in the JBP3-associated chromatin remodeling complex.

| Gene ID | Name | HHpred/InterPro domains | Tagged protein |  |  |
| --- | --- | --- | --- | --- | --- |
|  |  |  | JBP3 | J3C-Chromo | J3C-CS |
| LtaP36.0380 | JBP3 | SWI1, J-binding & N-terminal TFIIIS | 15.9 | 7.9 | 7.8 |
| LtaP35.2400 | SET-J3C | SET & C4-type Zn finger | 12.8 | 8.2 | 9.4 |
| LtaP14.0150 | Chromo-J3C | Chromo & chromo shadow | 11.7 | 11.6 | 8.8 |
| LtaP28.2640 | CS-J3C | Chromo shadow | 12.5 | 9.8 | 11.8 |
| LtaP12.0900 | HPC-J3C | None found | 12.1 | 9.8 | 6.3 |

Therefore, we have dubbed this protein Chromo-J3C and predict that it may be involved in recognition of methylated lysine residues on histone tails. LtaP28.2640 is annotated as a “Hypothetical protein, conserved”, but residues 187-227 also show structural similarity to the Chromo shadow domain (**Fig. S3F**), and so we have called it CS-J3C. The *LtaP12*.*0900* gene is misassembled in the *L. tarentolae* reference genome, so we used full-length orthologues from other *Leishmania* genomes for subsequent analyses. However, BlastP, InterProScan and HHpred analyses were uninformative, so we have called this protein HPC-J3C (for Hypothetical protein, conserved in J3C). Phylogenetic analysis showed that HPC-J3C has poor sequence conservation, even in other trypanosomatids, with orthologues in other genera being shorter than in *Leishmania*. HHpred analysis of the *T. brucei* orthologue (Tb927.1.4250) showed structural similarity at the C-terminus to subunits from several large protein complexes involved in a variety of processes, including histone remodeling. (**Fig S3G**).

To confirm association of these proteins with JBP3 (and each other), we transfected TAP-tagged versions of them into *L. tarentolae* T7-TR. Unfortunately, cloning of *HPC-J3C* failed because of errors in the genome sequence (see below) and transfectants containing the *SET-J3C* construct did not express the tagged protein (perhaps because over-expression was deleterious for cell growth), but the Chromo-J3C and CS-J3C transfectants expressed tagged proteins of the expected size (see **Fig. S1B**), enabling affinity purification (**Fig. 3A**). Subsequent mass spectrometric analysis of co-purifying proteins showed that Chromo-J3C and CS-J3C pull-downs enriched JBP3 and the same JBP3-associated proteins (**Table 2** and **Table S1**), indicating that they form a separate JBP3-containing complex. Bioinformatic analyses (described above) suggest that the proteins in this complex are likely associated with chromatin modification and/or remodeling, prompting us to call it the JBP3-associated chromatin complex (abbreviated as J3C).

**FIG 3.**
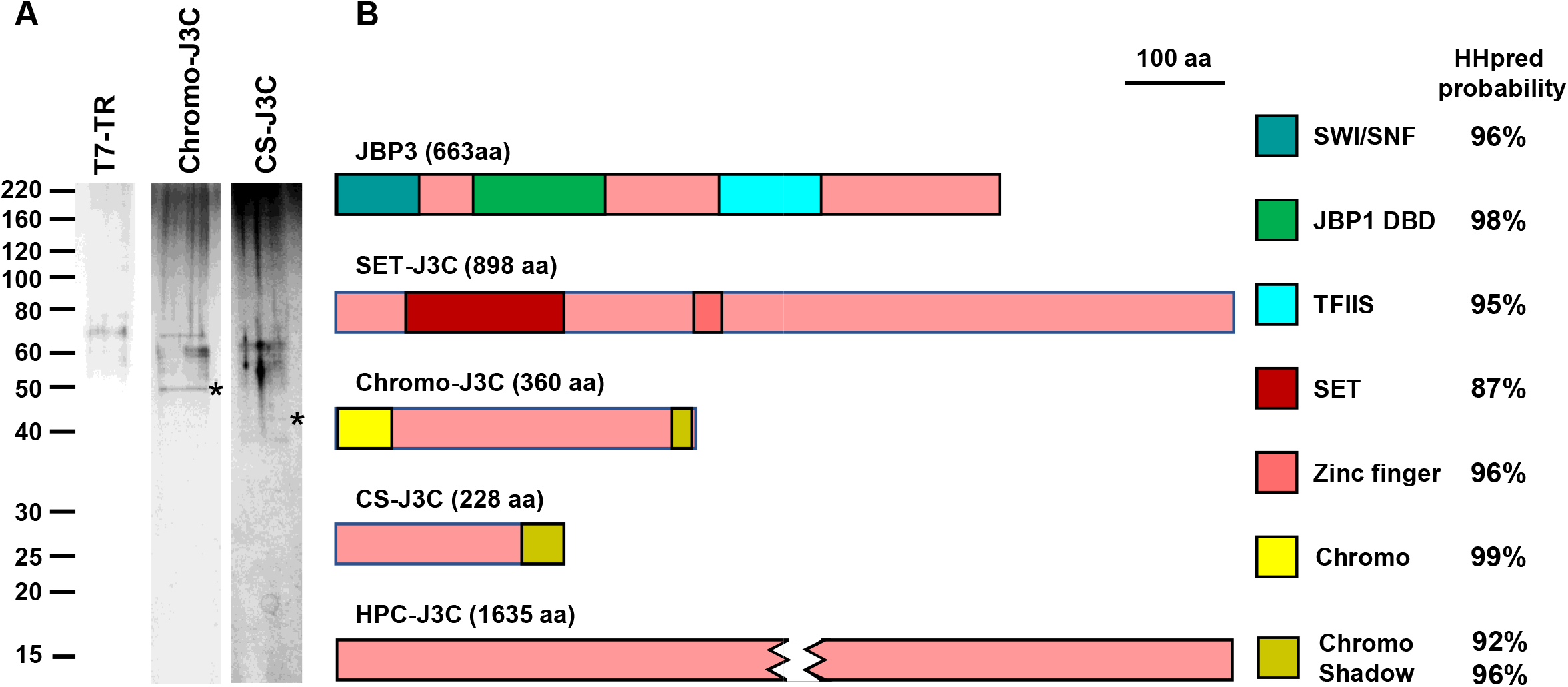
The JBP3-associated chromatin complex. **A**. Proteins that co-purify with TAP-tagged Chromo-J3C and CS-J3C over-expressed in T7-TR cells were analyzed by SDS-PAGE and silver staining as described in Fig. 1. **B**. Schematic representation showing the key domains of the five proteins that co-purified in the JBP3-associated chromatin (J3C) complex,

### JBP3 interacts with PAF1C

In addition to the components of the PJW/PP1 and J3C complexes, two other proteins (encoded by *LtaP35*.*2870* and *LtaP29*.*1270*) were substantially enriched (>110-fold and >23-fold respectively) in both JBP3 TAP-tag experiments (**Table 3** and **Table S1**). LtaP35.2870 is annotated in TriTrypDB as “RNA polymerase-associated protein LEO1, putative” and this homology was confirmed by HHpred analyses (**Fig. S3H**). LEO1 is a subunit of the RNAP II-associated factor 1 complex (PAF1C), which facilitates transcription elongation by regulating chromatin modification (35-37). Interestingly, mass spectrometric analysis of proteins that co-purified with TAP-tagged LEO1 (**Fig. 4A** and **Table 3**) did not detect JBP3, but identified three proteins (encoded by *LtaP36*.*4090, LtaP29*.*2750*, and *LtaP29*.*1270*,) that were enriched >630-fold in both experiments (**Table S1**). The first two were also enriched >50-fold in one of the two JBP3 TAP-tag experiments (**Table S1**) and are obvious homologues of PAF1C subunits. HHpred analyses showed that LtaP36.4090 contains the Ras-like fold characteristic of C-terminal domain of CDC73 subunit of PAF1C (**Fig. S3I**) and is annotated as such on TriTrypDB. LtaP29.2750 contains several tetratricopeptide repeat (TPR) domains implicated in protein-protein interactions and shows considerable overall similarity to the CTR9 subunit of PAF1C (**Fig. S3J**). Functional studies of the *T. brucei* CTR9 orthologue (Tb927.3.3220) indicated that it is essential for parasite survival and depletion of its mRNA reduced the expression of many genes involved in regulation of mRNA levels (38). The third protein (LtaP29.1270), which was also substantially enriched in both replicates of the TAP-tagged JBP3 experiments, is not an obvious orthologue of any known PAF1C subunit. This protein is annotated as a “Hypothetical protein, conserved” in TriTrypDB, but HHpred analysis (**Fig. S3K**) revealed a central domain (amino acids 232-358) with structural similarity to the PONY/DCUN1 domain found in DCN (Defective in Cullin Neddylation) proteins that are involved in regulation of ubiquitin ligation cascades (39). Our results (**Table 3**) suggest that LtaP29.1270, which we will refer to as DCNL (DCN-like) hereafter, forms an integral part (along with LEO1, CDC73 and CTRL) of a PAF1C-like (PAF1C-L) complex in *Leishmania*. The lack of significant reciprocal enrichment of JBP3 in the LEO1-TAP pull-downs (this paper) and its absence from the CTR9-TAP pull-downs in *T. brucei* (38) suggests a transient and/or indirect association of JBP3 with PAF1C-L.

**Table 3.** Enrichment of proteins in the PAF1C-L complex.

| Gene ID | Name | HHpred/InterPro domains | Tagged protein |  |
| --- | --- | --- | --- | --- |
|  |  |  | JBP3 | LEO1 |
| LtaP35.2870 | LEO1 | Leo1-like protein | 7.4 | 16.8 |
| LtaP29.1270 | DCNL | PONY/DCUN1 domain | 5.4 | 13.9 |
| LtaP36.4090 | CDC73 | Cell division control protein 73 (C-terminal) | (5.7) | 10.4 |
| LtaP29.2750 | CTR9 | RNA polymerase-associated protein Ctr9 | (6.2) | 9.5 |
| LtaP14.0860 | RTF1L | Plus-3 domain | 0.6 | 3.8 |

Parentheses indicate that enrichment was observed in only one replicate.

**FIG 4.**
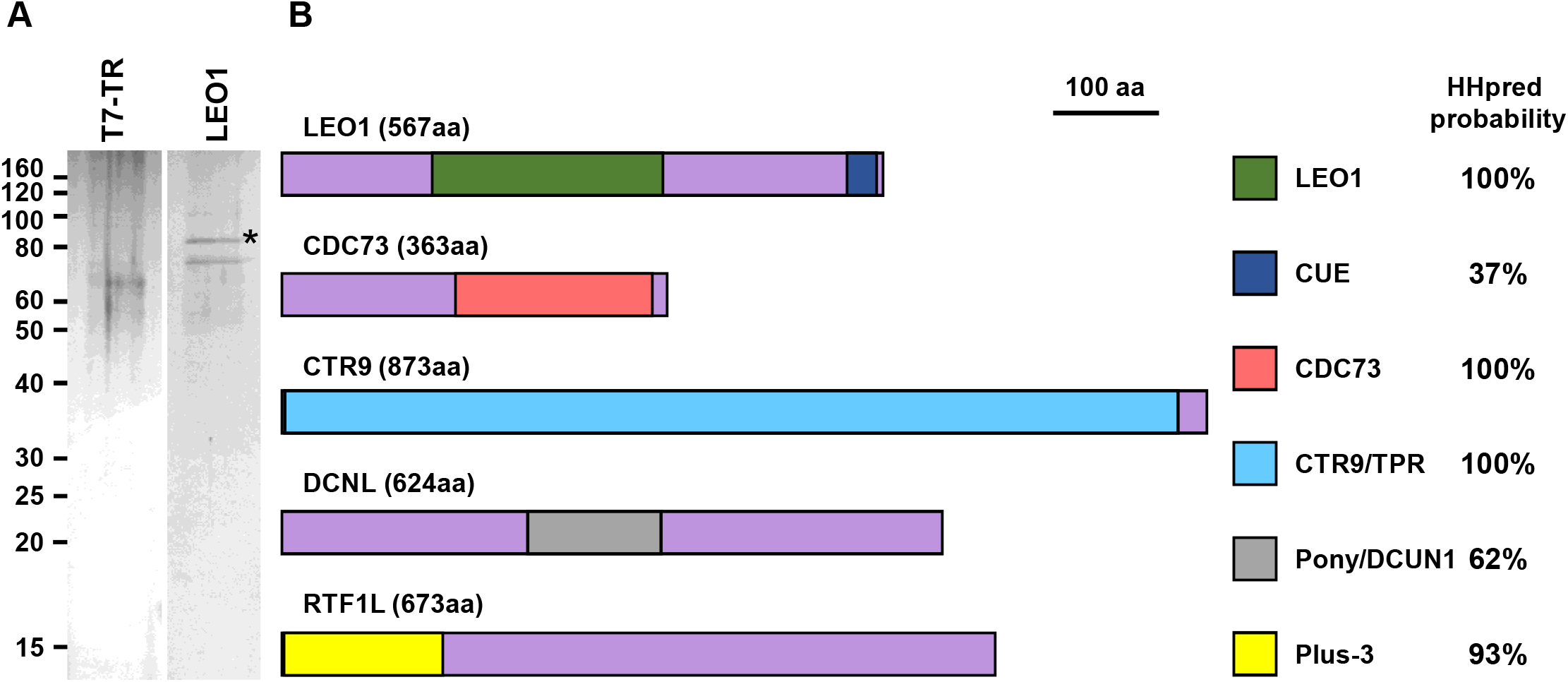
The PAF1C-like complex. **A**. Proteins that co-purified with TAP-tagged LEO1 overexpressed in T7-TR cells were analyzed by SDS-PAGE and silver-staining as described in Fig. 1. **B**. Schematic representation showing the key domains of five components of the PAF1C-L complex and their probability score from HHpred analysis.

TAP-tagging of *T. brucei* CTR9 by others (38) revealed the same constellation of PAF1C-L subunits (LEO1, CDC73 and DCNL), as well an additional protein (Tb927.7.4030). Close examination of our results revealed that the *Leishmania* orthologue (LtaP14.0860) of Tb927.7.4030 is also enriched >10-fold in both LEO1 TAP-tag experiments (**Table S1** and **Table 3**). While this protein is annotated “Hypothetical protein, conserved” in TriTrypDB, HHpred analysis revealed an N-terminal (amino acids 2-152) structural similarity to the Plus-3 domain of human RTF1 (**Fig. S3L**), a component of human and yeast PAF1C. Thus, LtaP14.0860 (which we have dubbed RTF1L) is likely the functional equivalent of RTF1, although, in *Leishmania*, it may be less tightly associated with PAF1C-L.

### Depletion of JBP3 causes defects in transcription termination

The results of the TAP-tag experiments presented above suggest that JBP3 is an integral part of two protein complexes (PJW/PP1 and J3C) and interacts, possibly indirectly, with another (PAF1C-L). Since similar complexes have been associated with chromatin modification/remodeling and regulation of transcription in other organisms, we postulated that JBP3 may mediate transcription termination in *Leishmania*. To test this hypothesis, we used CRISPR/Cas9 (40) to delete *JBP3* in the *L. tarentolae* cell-line bearing a tetracycline (Tet)-regulated copy of *JBP3-TAP* (see above). We were able to delete both endogenous copies of *JBP3* only when *JBP3-TAP* expression was induced with Tet, suggesting that it is essential in *Leishmania*. To interrogate the effect of JBP3 depletion, we grew two stable transfectants lacking both endogenous copies of *JBP3* (but containing a Tet-regulated copy of *JBP3-TAP*) for 8-11 days in the presence or absence of tetracycline **(Fig. 5A)**. While cells grown in the presence of drug maintained a constant growth rate (with a generation time of ∼9 hours) over the length of the experiment; JBP3-TAP protein levels decreased markedly during the first day after removal of tetracycline, dropping to ∼2% of the initial level by day 2 (**Fig. 5B**). The growth rate in the absence of drug slowed after day 3, with the generation time increasing to >20 hours on day 6, before returning to almost the parental wild-type rate after day 10 (**Fig. 5A**).

**FIG 5.**
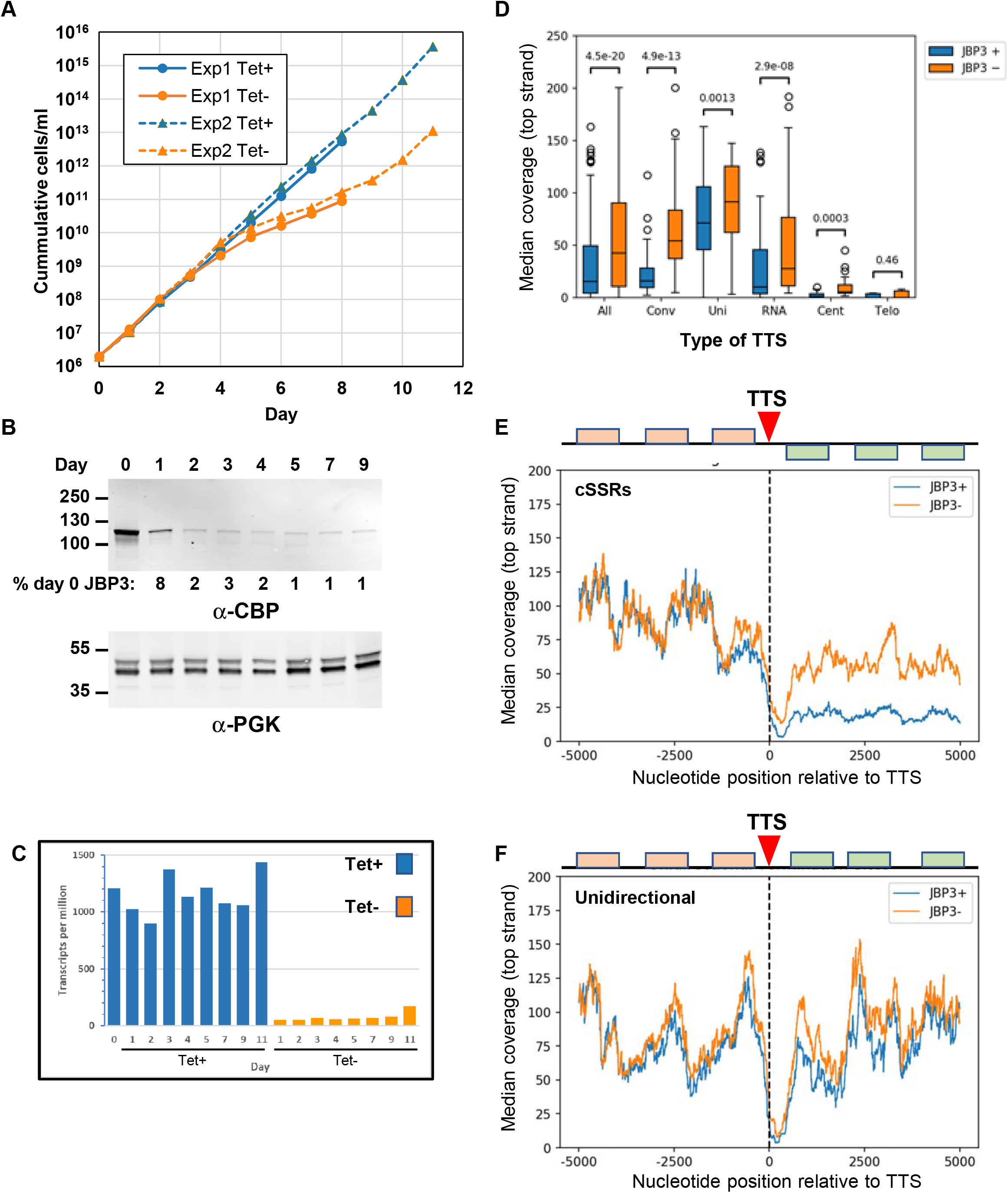
Depletion of JPB3 results in read through at transcription termination sites. **A**. Growth analysis. Two independently generated *L. tarentolae* clones lacking endogenous *JBP3* but containing a Tet-regulated TAP-tagged *JBP3* gene was grown in the presence (blue lines) or absence (orange lines) of Tet. The numbers on y-axis are corrected for dilution during sub-culturing. The solid lines show a clone where the *JBP3* genes were replaced by *pac* (and grown in the presence of puromycin), while the dotted lines show a clone where with the *JBP3* genes replaced by *neo* (and grown in G418). **B**. The level of TAP-tagged JBP3 expressed by the T7-TR/JBP3-MHTAP/Δ*jbp3*::*neo* clone grown in the absence of Tet was monitored by Western blot analysis using antibodies against the calmodulin binding peptide (CBP) of the TAP-tag. Antibodies against phosphoglycerate kinase (PGK) served as a loading control. The percent JBP3 levels in comparison to day 0 is shown below the anti-CBD blot. **C**. *JBP3-TAP* mRNA levels for the T7-TR/JB3-MHTAP/Δ*jbp3*::*neo* clone grown in the presence (blue) or absence (orange) or Tet for the number of days indicated. mRNA levels are expressed transcripts per million (TPM) as determined by RNA-seq analysis using Geneious. **D**. Box-and-whiskers plots showing the median top strand coverage in the 5 kb downstream of all 192 TTSs (All). Separate plots are shown for the 46 TTS at cSSRs (Conv), 30 TTSs between head-to-tail PTUS (Uni), 39 TTS immediately upstream of one or more RNA genes (RNA), 21 TTS adjacent to a centromere (Cent) and 56 TTSs at telomeres (Telo). **E**. Median of top strand coverage at each nucleotide position in the 10 kb surrounding the 46 TTS at cSSRs. The schematic represents the protein-coding genes associated with each strand at an “average” cTTS. The second PTU at each cSSR is re-oriented so that the genes are represented on the top strand. **F**. Median of top strand coverage at each nucleotide position in the 10 kb surrounding the 30 TTS at between unidirectional (head-to-tail) PTUs. The schematic represents the protein-coding genes associated with each strand at an “average” uTTS.

To assess the role of JBP3 on mRNA expression and transcription termination, RNA was isolated daily from the cells after Tet withdrawal and used to generate strand-specific RNA-seq libraries. Illumina sequencing reads were mapped to the *L. tarentolae* reference genome and normalized read counts calculated for every gene. Differential expression analysis revealed that *JBP3-TAP* mRNA levels were ∼20-fold lower in the absence of Tet **(Fig. 5C**). Interestingly, there was a marked increase in the *JBP3-TAP* mRNA levels in the Day 11 Tet-sample, coincident with resumption of normal growth. Consequently, this sample was excluded from subsequent analyses, along with the Day 1 Tet-samples (since JBP3 protein was still ∼8% of the initial level). Further analysis (using the DESeq2 module of Geneious) revealed that 17 genes had significantly higher mRNA abundance (>2-fold, p<0.001) in the remaining Tet-samples, which all had low JBP3 protein levels (**Table S2**). Interestingly, 14 of these genes are located adjacent (or close) to transcription termination sites (TTSs). Indeed, 34 of the 50 most up-regulated genes are located near TTSs, with 24 of these located at convergent strand-switch regions (cSSRs) where the 3’ termini of two PTUs converge.

To further characterize these increases in RNA abundance, we analyzed the read coverage for 5 kb on either side of all 192 TTSs in the *L. tarentolae* genome. As expected, when JBP3 is expressed (Tet+ samples) the median-normalized coverage on the top (coding) strand decreased sharply downstream of the TTS (**Fig. S4A**). However, in samples with very low JBP3 protein levels (days 2-9 Tet-), the read coverage downstream of the TTS was significantly higher, suggesting that reduction of JBP3 resulted in substantial transcriptional read-through. Importantly, this increase in read-through transcription did not occur to the same extent at different types of TTS (**Fig. 5D**). It was most pronounced at the 23 non-centromeric cSSRs without RNA genes (**Fig. 5E** and **Fig. S4B**), where transcript abundance was almost as high downstream of the TTS as it was upstream. There was also a significant increase in read-though transcription downstream of the TTS between unidirectionally (head-to-tail) oriented PTUs (**Fig. 5F** and **Fig. S4C**); although it was more subtle, since the gap between each PTU is small. Conversely, there was only a small increase in read-through at TTSs upstream of RNA genes transcribed by RNAP III (**Fig. S4D**), and essentially no read-through at centromeres (**Fig. S4E**) or telomeres (**Fig. S4F**). Analysis of bottom (non-coding) strand transcripts revealed no significant differences between Tet+ and Tet-samples (left-hand panels in **Fig. S4**), except at cSSRs, where read-through from the second PTU results in a substantial increase in antisense transcripts upstream of the TTS (**Fig. S4B**), due to read-through from the convergent downstream PTU. Similar analysis of transcript abundance surrounding transcription start sites (TSSs), revealed no significant changes due to JBP3 depletion, except for a small increase in top (coding) strand coverage when PTUs were oriented unidirectionally (**Fig. S5C** and **S5G**), presumably due to read-through from the preceding PTU. Importantly, there was little or no increase in bottom (non-coding) strand coverage upstream of most TSSs.

## Discussion

Using the trypanosomatid-specific GT, which carries out the second step of J biosynthesis, as an entrée to search for the molecular machinery associated with regulation of transcription in *Leishmania*, we have identified a network of three protein complexes that contain conserved building blocks often used to assemble molecular machinery regulating transcription in other eukaryotes. A novel J-binding protein (JBP3) lies at the nexus of these complexes (**Fig. 6**) and provides new insight into the molecular mechanism(s) used to mediate transcription termination at the end of the polycistronic transcription units emblematic of these (and related trypanosomatid) parasites. We have shown that JBP3 plays a central role in controlling termination of RNAP II transcription, since depletion of JBP3 leads to defects in transcriptional termination at the 3’ end of PTUs in *Leishmania* (**Fig. 5**), just as it does in *T. brucei (20)*. However, read-through transcription is not seen to the same extent at all TTSs. The presence of RNAP III-transcribed RNA genes downstream of the TTS appears to effectively block RNAP II, as we have seen previously for JBP2 null mutants (11), and there is little or no read-through at TTSs immediately upstream of centromeres and telomeres. This suggests that factors other than JBP3 may play a role in reducing transcriptional read-through at these loci. Alternatively, it is possible that the higher J content at centromeres and telomeres may more effectively “capture” what little JBP3 remains in the Tet-cells. In contrast to the recent results from *T. brucei* (20),we find little evidence for antisense transcription at the 5’ end of PTUs in *Leishmania*, even after depletion of JBP3 levels. Divergent strand-switch regions (dSSRs), where adjacent PTUs are on opposite strands, tend to be smaller in *Leishmania* than those in *T. brucei* and lack the J-containing DNA found in the latter, suggesting that there may be little inappropriate antisense transcription in these regions. However, we do find evidence that depletion of JBP3 in *Leishmania* results in up-regulation of mRNA levels for protein-coding genes at the 3’ end of PTUs (**Table S2**). It is possible that this phenomenon is due to more efficient polyadenylation of transcripts caused by uncovering of cryptic trans-splicing sites downstream of the normal TTS. Toxic effects of a gradual accumulation of proteins from these mRNAs may be one explanation for the lag between appearance of defects in transcription termination (day 2) and decrease in growth rate (day 4). This lag period also indicates that read-through transcription if not an artifact of reduced growth rate.

**FIG 6.**
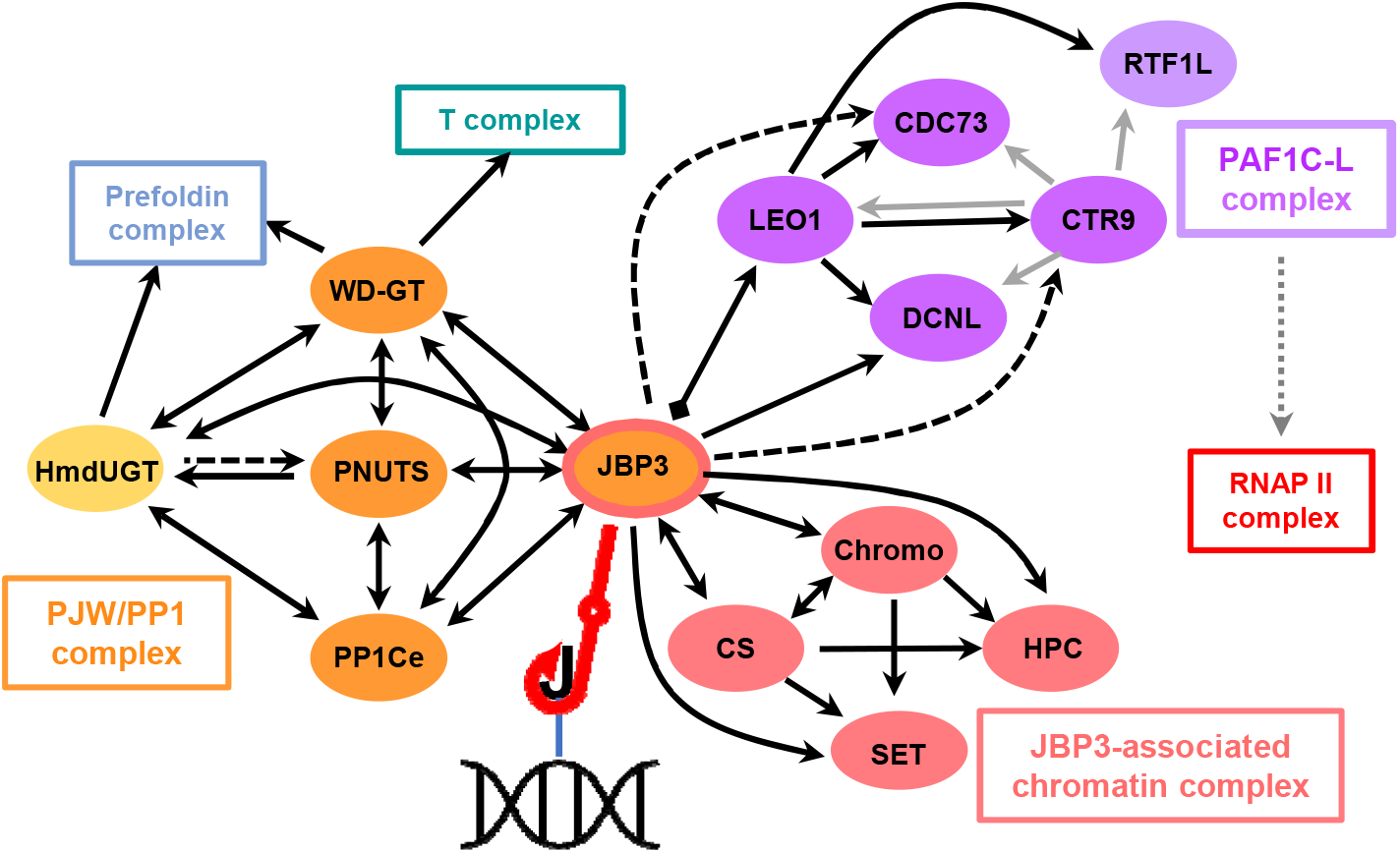
Network of interactions between JBP3-assocated protein complexes. Solid lines denote proteins enriched in both replicates of the TAP-tag pull-downs and dashed lines indicate proteins enriched in only one sample. Double headed lines represent reciprocal enrichment with both proteins used as bait, while a diamond shape at one end of the line indicates that JBP3 was not detected in LEO1 pull-down. Lines with a single arrowhead indicate that reciprocal enrichment was not attempted. Grey arrows represent interactions identified by co-purification of proteins with CTR9 in *T. brucei* (38). Subunits within the three distinct JBP3-associated protein complexes are denoted by different colors, while the chaperone and RNA polymerase complexes are represented by boxes without individual components. The dotted line connecting the PAFC1-L complex to RNAP II reflects the interactions observed in other organisms. The interaction between JBP3 and base J is marked by the red fishhook.

Our initial TAP-tagging experiments showed that GT associates (directly or indirectly) with four other proteins that resemble components of the PTW/PP1 complex, which PNUTS, TOX4, WDR82 and PP1C) has been implicated in numerous different cellular processes; including control of chromatin structure during cell cycle progression (27), repair of DNA damage by non-homologous end-joining (41), maintenance of telomere length (41), and developmental regulation of transcription (42). The *Leishmania* PJW/PP1 complex shows obvious parallels to the metazoan PTW/PP1 complex by incorporating analogous (although not necessarily homologous) proteins, with JBP3 substituting for the DNA binding function of TOX4. However, there are some interesting differences between *Leishmania* PJW/PP1 and the homologous *T. brucei* complex; most notably the absence of PP1C in the latter, where the complex was therefore called PJW (20). This absence is intriguing in the light of a recent publication that showed the mammalian PTW/PP1 complex dephosphorylates the transcription elongation factor Spt5, thereby causing the RNAP II transcription complex to decelerate within the termination zone downstream of poly(A) sites and allowing the Xrn2 exonuclease to “track down and dislodge” the polymerase from the DNA template (30). This suggests that *Leishmania* and American trypanosomes may use PJW/PP1 to dephosphorylate Spt5 and mediate transcription termination, but that African trypanosomes do not use this mechanism. In addition, although *T. brucei* encodes an orthologue of GT, it was not found to be associated with the PJW complex, pointing to another potential biological difference from *Leishmania* (or perhaps merely reflecting our use of a more rapid and sensitive purification protocol). In mammalian PTW/PP1C, PNUTS not only contains the phosphatase inhibitor motif, but also contains a nuclear localization signal (NLS) and provides a “scaffold” for recruiting the other proteins. However, *Leishmania* PNUTS is much smaller and lacks an obvious NLS, so it is possible that other proteins in the complex provide these functions. For example, JBP3 contains a domain with structural similarity to the N-terminal TFIIS protein interaction domain found in mammalian PNUTS and WD proteins can act as a scaffold in other complexes (43, 44).

JBP3 is also present in a complex (J3C) that contains proteins with domains suggesting a role in chromatin modification and/or remodeling. One protein (SET-J3C) contains a SET domain, which is often present in histone methyltransferases (HMTs) that modify lysine and arginine residues in histone proteins. HMTs usually also contain cysteine-rich pre-SET and post-SET domains that play a crucial role in substrate recognition and enzymatic activity by coordinating zinc ions. SET-J3C contains a Zn finger domain downstream of the SET domain (Fig. S3D) that may fulfill similar functions, suggesting it is likely to have HMT activity. Two other proteins (Chromo-J3C and CS-J3C) contain Chromo and/or Chromo shadow domains typically involved in recognition of the methylated lysine residues on histone tails and may be functional homologues of the metazoan heterochromatin protein 1 (HP1) and/or fission yeast Swi6, which contain similar domains and are involved in repression of gene expression by heterochromatin (34). Thus, it is possible that J3C facilitates re-packing of chromatin at J-containing regions of the genome, rendering them heterochromatin-like and inaccessible to RNA polymerases. The function of the HPC-J3C subunit is unknown at this time, but the *T. brucei* orthologue (encoded by *Tb927*.*1*.*4250*) shows structural similarity to subunits from different protein complexes implicated in a several processes, including histone modification and/or chromatin remodeling, and it localizes to the nucleus (45). This suggest that the primary function of HPC-J3C may be to serve as a scaffold for recruitment and/or stabilization of the J3C complex.

JBP3 also associates (albeit transiently and/or indirectly) with a third protein complex (PAF1C-L), providing another potential connection between J and regulation of transcription. PAF1C-L contains proteins with functional domains similar to those in yeast and mammalian PAF1C, which associates with the large subunit of RNAP II (35, 46) and plays a critical role in transcription elongation and termination (47, 48). Four proteins (PAF1, CDC73, LEO1 and CTR9) are consistently found as components of PAF1C in all other eukaryotes, while RTF1 (49) and the WD40-containing protein Ski8/WDR61 (50) show a less ubiquitous association. The parallels between the mammalian and trypanosomatid complexes are obvious, since homologues of the CDC73, LEO1 and CTR9 subunits copurify with JBP3 and experiments performed by others in *T. brucei* showed that CTR9 has a tight association with LEO and CDC73 (38). Our TAP-tag experiments using LEO1 as the bait protein showed enrichment of CDC73 and CTR9, along with a protein containing a Plus-3 domain similar to that in RTF1. Interestingly, the Plus-3 domain of RTF1 has been implicated in binding to Spt5 (51), which is tantalizing in light of a potential role for the PJW/PP1 complex in dephosphorylating this transcription factor (30). We (and others) failed to identify a convincing homologue of Ski8/WDR61 (or any other WD40 protein) in the PAF1C-L complex, but this protein is not tightly associated with mammalian PAF1C either. Remarkably, the namesake of the complex, PAF1, is absent from pull-downs of LEO and CTR9 in *L. tarentolae* and *T. brucei* respectively, even though it (along with CTR9) is essential for assembly of the complex in both yeast and humans (52). Moreover, extensive bioinformatic analysis of the trypanosomatid genomes failed to identify a homolog of PAF1, suggesting that is truly absent from *Leishmania* and *Trypanosoma* PAF1C-L complex. However, PAF1C-L contains an additional, trypanosomatid-specific, component (DCNL) that has a putative protein-binding domain with structural similarity to the PONY/DCUN1 domain found in the eukaryotic DCN protein family. In other eukaryotes, DCN1 is required for neddylation of cullin in SCF-type E3 ubiquitin ligase complexes that mark cellular proteins for proteosomal degradation (53). It is interesting to speculate that DCNL may be involved in PAF1C-L recruitment/function by interaction with N-terminal acetylated residues on histones and/or other chromatin-associated proteins. Whether DCNL functionally replaces PAF1 will remain an open question until its molecular function is dissected in more detail.

We have previously postulated that J might terminate transcription by directly preventing progression of polymerase (11). The data presented here suggest three alternative, but not mutually exclusive, hypotheses based on the ability of JBP3 to bind base J. Firstly, JBP3 may recruit the PJW/PP1 complex to J-containing regions, where it helps stall the elongating RNAP II by dephosphorylating Spt5 and causing the transcription complex to decelerate within the termination zone. Secondly, J3C complex may enhance termination by modifying and/or remodeling the chromatin at J-containing regions, thereby preventing passage of RNAP II. Thirdly, we speculate that the interaction between JBP3 and PAF1C-L may also tether the RNAP II to the termination zone, where the DCNL subunit ubiquitinates the polymerase complex, promoting its degradation by the proteasome (54, 55). While disruption of the PJW complex in *T. brucei* (20) provides additional evidence for the first hypothesis, it remains to be seen whether disruption of J3C and PAF1C-L complexes also cause a defect in transcription termination.

While our findings provide several novel insights into the role of base J in transcription termination, they also raise several interesting questions. For example, why is GT part of the PJW/PP1 complex? One could envisage that recruitment of the PJW/PP1 complex to regions of the genome containing J recruits may allow more efficient glucosylation of nearby HOMedU residues. Our proteomics data suggest that JBP3 interacts to varying degrees with three different protein complexes (J3C, PJW/PP1 and PAF1C-L). It will be important to understand how these interactions are regulated. Are the complexes present at the same region simultaneously or are they temporally and/or spatially segregated? For example, PJW/PP1 may be present at the ends of all PTUs, while J3C is present only at centromeres and/or centromeres. What histone modification/remodeling is mediated by J3C and what role do this play a role in transcription termination? The availability of modern genome-wide approaches will no doubt provide the appropriate tools to answer these questions.

## Materials and methods

### Plasmid construction

To create an expression vector that expresses epitope-tagged transgenes in *Leishmania*, the MHTAP tag (which bears a Myc epitope, six histidines, a protein A domain, and calmodulin binding peptide) was amplified from the plasmid pLEW-MHTAP (56) with the primers MHTAP-BamHI-S and MHTAP-Not1-AS (all primers use in this study are described in **Table S3**). Following cleavage with BamHI and NotI, the PCR fragment was inserted into BglII+Not1-digested pLEXSY-I-bleCherry3 (Jena Biosciences). The resulting plasmid (pLEXSY-MHTAP) allows the TAP-tagging of introduced coding regions under the control of a Tet-regulated T7 promoter, and insertion into the *ODC* locus on chromosome 12 of *L. tarentolae*. We used a combination of published datasets to identify the 5’ end of *L. tarentolae* mRNAs as marked by the 39 base pair splice leader sequence (11) and ribosome profiling data from *L. donovani* (unpublished data), to identify the correct CDS for bait proteins. CDSs were PCR amplified and digested with the restriction enzymes indicated in **Table S3** prior to cloning.

### Parasite strains and tissue culture

The *Leishmania tarentolae* Parrot-TarII wild-type (WT) and T7-TR strains (Jena Bioscience) were grown in SDM-79 medium supplemented with 10% fetal bovine serum. Strain T7-TR has constitutively expressed T7 RNA polymerase and Tet repressor genes integrated into the rDNA locus, allowing for Tet-induced expression of integrated (or ectopically expressed) genes. Nourseothricin and hygromycin B were added to the medium at 100 µM to maintain expression of T7 RNA polymerase and Tet repressor (TetR).

### Tandem affinity purification of tagged protein complexes

Ten µg of *Swa*I-digested pLEXSY-MHTAP plasmid encoding a TAP-tagged protein was electroporated into the *L. tarentolae* wild type (WT) and T7-TR cell-lines as described (57) and transfectants selected with 100 µg/ml bleomycin and maintained in 20 µg/ml bleomycin (plus nourseothricin and hygromycin B as described above). Proteins associated with the TAP-tagged “bait” were purified from 500 ml of cells following overnight culture (in the presence of 2 µg/ml tetracycline for T7-TR transfectants) as described (56), except that NP-40 was omitted from the final four washes of the proteins on the calmodulin beads and from the calmodulin elution buffer. The protocol went from lysis of cells to purified samples within 6 hours. A sample from each pull-down (5% of the total eluate) was separated by 4-20% SDS-PAGE and proteins were visualized using SilverQuest Stain (Thermo Fisher Scientific Life Technologies). Fractions containing a protein with the predicted molecular weight of the bait (usually fractions 2 and 3) were pooled. The same fractions were pooled from mock TAP purifications of the control parental line not expressing any bait protein.

### Western blotting

Proteins from transfected cells were separated by SDS-PAGE on 4-20% gradient gels, transferred onto nitrocellulose and detected with either mouse anti-6×His (Clontech) at 0.25 µg/ml or rabbit antibody against calmodulin binding peptide Calmodulin Binding Peptide (GenScript) at 0.1 µg/ml, with rabbit antibody against *T. brucei* phosphoglycerate kinase serving as a control (58). Primary antibodies were detected with goat anti-rabbit Ig conjugated with AlexaFluor 680 (50 ng/ml) or goat anti-mouse Ig conjugated with IRDye 800 (25 ng/ml) and imaged on the LI-COR Odyssey CLX.

### Proteomic analysis

Pooled protein fractions were denatured with 6 M urea, reduced with 5 mM dithiothreitol, alkylated with 25mM iodoacetamide, and digested at 37°C for three hours using 1:200 w:w endoproteinase Lys-C (Thermo Fisher Scientific). The urea was then diluted to 1.5 M and samples further digested at 37°C overnight with 1:25 w:w trypsin (Thermo Fisher Scientific). Proteinase activity was stopped with formic acid, and peptides purified using C18 reversed-phase chromatography (Waters), followed by hydrophilic interaction chromatography (HILIC; Nest Group). Purified peptides were separated by online nanoscale HPLC (EASY-nLC II; Proxeon) with a C18 reversed-phase column (Magic C18 AQ 5um 100A) over an increasing 90 minute gradient of 5-35% Buffer B (100% acetonitrile, 0.1% formic acid) at a flow rate of 300 nl/min. Eluted peptides were analyzed with an Orbitrap Elite mass spectrometer (Thermo Fisher Scientific) operated in data-dependent mode, with the 15 most intense ions per MS1 survey scan selected for MS2 fragmentation by rapid collision-induced dissociation (rCID) (59). MS1 survey scans were performed in the Orbitrap at a resolution of 240,000 at m/z 400 with charge state rejection enabled, while rCID MS2 was performed in the dual linear ion trap with a minimum signal of 1000. Dynamic exclusion was set to 15 seconds.

Raw output data files were analyzed using Maxquant (v1.5.3.30) (60) to search against a proteome predicted after resequencing and annotation of the *L. tarentolae* Parrot (LtaP) genome (Sur *et al*, manuscript in preparation). A reverse sequence decoy database was used to impose a strict 1% FDR cutoff. Label-free quantification was performed using the MaxLFQ algorithm (61) and further data processing was performed in Perseus (v1.5.3.1) (62) and Microsoft Excel. To avoid zero-value denominators, null values in the remaining data were replaced by imputation using background signal within one experiment using Perseus. Non-parasite contaminants, decoys, and single peptide identifications among all samples in an experiment were removed. Proteins were deemed to be part of a complex associated with the bait protein if at least two peptides were detected and the protein showed more than 32-fold (log_2_ >5) enrichment (compared to the control not expressing the bait) in both replicates. In a few cases, proteins showing >100-fold enrichment in a single replicate only, were also considered as potential subunits of the complex. Proteins with 1024-fold (log2 <10) less enrichment than the bait protein were assumed to be co-purifying contaminants and (usually) ignored. Mass spectrometry data was deposited in the MassIVE database (https://massive.ucsd.edu/ProteoSAFe/static/massive.jsp) and can be accessed from ProteomeXchange using the identifier PXD020779.

### Bioinformatic analysis of protein function

Structure-based similarity searches for known domains were performed with HHPRED (*23*). Domain boundaries for JBP3 DNA binding domain (DBD) were refined by aligning trypanosomatid sequences that clustered in the same OrthoMCL group as JBP3 with T-Coffee (63). Homology models of the JBP3-DBD domain using RosettaCM (64) were built with the *L. tarentolae* JBP1 structure (PDB: 2XSE) as a template. The top-scoring model covered residues 111 to 312 of JBP3-DBD with a confidence score of 0.67.

### Deletion of JBP3 using SaCas9

JBP3 was deleted using *Staphylococcus aureus* Cas9 (SaCas9)-directed cleavage of sites flanking the endogenous locus as described (40). Briefly, guide RNAs directed at sites for SaCas9 cleavage were generated *in vitro* using T7 Megashort from ThermoFisher from PCR-generated templates. The 5’ and 3’ gRNA sequences used were GATGTGAAACGCTAAGCAGTC<u>CCGAGT</u> and AGGAACGAAAGCACACAGCAG<u>AGGAGT</u>, where the PAM sites are underlined. Repair fragments containing a drug resistance gene were generated as described (65) from pTNeo or pTPuro templates using primers LtJBP3-up and LtJBP3-down. Heat-denatured guide RNA was complexed with 20 µg SaCas9 recombinant protein at equimolar ratio and incubated for 15 minutes at room temperature before mixing with 2 ug of each repair fragment that had been ethanol-precipitated and resuspend in Tb-BSF buffer (66). *L. tarentolae* T7/TR cells were grown overnight (in the presence of 2 ug/ml tetracycline for the latter), pelleted, washed with PBS, and resuspended in 100 µl Tb-BSF buffer. For each transfection, 10^6^ cells were mixed with the SaCas9/guide RNA complexes and repair fragments and electroporated in an Amaxa Nucleofector using program X-001. Immediately following transfection cells were split into three flasks. After allowing the cells to recover overnight, antibiotics were added to the flasks; one of which was grown in 10 µg/ml G418, one in 4 µg/ml puromycin, and one with both drugs. Deletion of the endogenous *JBP3* gene(s), was confirmed by PCR amplification of genomic DNA using primers JBP3-M84P and JBP3-P2147M that flank the region being deleted. Clones cell lines were obtained by limiting dilution of the transfectants and clones were retested for JBP3 deletion by PCR. While we were able to obtain clones where both endogenous copies of *JBP3* were deleted by selection with either puromycin or neomycin, we were unable to obtains lines where both drugs were used.

### RNA-seq analysis

RNA was isolated using TRIzol (Thermo Fisher Scientific)) and resuspended in 10mM Tris, pH 7and RNA quality assessed using the Bioanalyzer 6000 Pico Chip (Agilent). mRNA was isolated from 1 µg total RNA using the NEB Poly(A) mRNA Magnetic Isolation Module (NEB) and prepared using the Stranded RNA-seq protocol (67), modified for *Leishmania* as described (*68*). Libraries were sequenced on an Illumina HiSeq, obtaining Paired End 150-bp reads.⁥⁥Reads were aligned against our in-house *L. tarentolae* genome with Bowtie2 (69) using the “very high sensitivity” parameter or the Geneious assembler (Geneious Prime 11.05, https://www.geneious.com) using the “Low Sensitivity/Fastest” option. Differential expression analysis was performed on the Geneious assemblies using the DESeq2 module to compare Tet+ and Tet-samples from days 2, 3, 4, 6 and 8 for replicate 1 and days 2, 3, 4, 5, 7 and 9 for replicate 2. Strand-specific read coverage was calculated directly from BAM files of the Bowtie2 alignments using customized pysam scripts (https://github.com/pysam-developers/pysam). RNA-seq data was deposited in the Sequence Read Archive (SRA) at NCBI and can be accessed using the BioProject Identifier <u>PRJNA657890</u>.

## Acknowledgements

We thank Lisa Jones of the Fred Hutchison Cancer Research Center Proteomics Resources facility for assistance with performing LC-MS/MS analysis. This work was supported in part by PHS grant R01 AI103858 (to P.J.M.) and contract HHSN272201700059C (to P.J.M.) from the National Institute of Allergy and Infectious Diseases.

## Author contributions

Concept and experimental design were done by B.C.J and P.J.M. Experiments were performed by B.C.J and J.R.M. Proteomic analysis was done by M.A.G. and J.A.R. Protein modeling and sequence analysis was done by I.Q.P. Analysis of RNA-seq data was done by A.S and P.J.M. Manuscript was written by B.C.J., M.P. and P.J.M. M.P. cracked the whip. All authors have reviewed the manuscript.

## Supplemental Files

**TABLE S1**

**Proteins that co-purify with TAP-tagged bait proteins**.

Only proteins with an average enrichment >32-fold (>64-fold for PP1Ce and 100-fold for WD-GT) and least two peptides in both experiments are shown, except for PNUTS and PFDN2 in the HmduGT pull-downs, CTR9 and CDC73 in JBP3 pull-downs, and RTF1L in the LEO1 pull-downs. The “bait” in HmdUGT-TAP Rep 1 was constitutively expressed (in wild type cells), while the bait in all other experiments was over-expressed by addition of tetracycline to T7-TR cell lines. The values for HPC-J3C represent the total number of peptides detected and the average fold-enrichment for 7 different CDSs for this gene, which was misassembled in the LtaP genome. Color shading is used to indicate proteins in the three JBP3-associated complexes described in **Fig. 6**. The lighter shading of HmdUGT indicates more transient association with the PJW/PP1 complex.

**TABLE S2**

**Genes with increased mRNA levels after depletion of JBP3**.

The 50 genes showing the largest average (between replicate 1 and replicate) significant (p<0.001) increase in RNA abundance, as determined by RNA-seq analysis, are shown in decreasing of log2 fold-change (FC). Different shading intensities indicate those genes increasing by more than 4-fold (dark), 2-fold (medium), or 1.5-fold (light). The “locus type” indicates whether the genes were immediately adjacent to (or within 20 kb of) convergent, unidirectional or telomeric transcription termination sites (cTTS, uTTS, and tTTS, respectively) or located within the central region of a polycistronic transcription unit (PTU-internal). TTSs located at centromeres are indicated by “-Cen”.

**TABLE S3**

**Oligonucleotide primers used for construct creation**.

Primer names include the restriction sites (underlined bases) used for cloning into pLEXSY-MHTAP. Bases preceding the restriction site were added to facilitate restriction digest of the PCR fragments. For the primers used for generate fragments for CRISPR/Cas9 deletion, the bases in bold match sequence flanking the JBP3 gene.

**FIG S1**

**Expression of proteins tagged with MHTAP**.

**A**. HmudGT tagged with MHTAP was transfected into either wild-type *L. tarentolae* Parrot-TarII or into the *L. tarentolae* strain T7-TR that expresses both T7 RNA polymerase and Tet repressor. Cell lysates were made from transfectants of the wild-type (WT) or T7-TR strain grown in the presence (+) or absence (-) of tetracycline (Tet). In the absence of Tet, the transgene in T7-TR is repressed, while in the presence it is induced. Protein from 5×10^6^ cells were separate by SDS-PAGE and transferred to a membrane and the tagged protein detected with α-CBP, which detects the calmodulin binding domain that is part of MHTAP.

**B**. Constructs expressing MHTAP tagged protein were transfected into T7-TR cells and lysates generated after treatment with Tet to induce expression of the transgene. Protein from 5×10^6^ cells were resolved by SDS-PAGE, transferred to a membrane and MHTAP-tagged protein detected with α-CBP.

**FIG S2**

**Phylogenetic tree for PP1C proteins from selected kinetoplastid species in comparison to proteins from human and *saccharomyces cerevisiae***.

Sequences were aligned using Muscle and viewed in Geneious (Geneious Prime 11.0.5, https://www.geneious.com). Genetic distances were calculated using Jukes-Cantor and trees constructed using neighbor-joining. The five clades of PP1C are shown, with genes from *Leishmania* (green dot), *T. brucei* (blue dot), *T. cruzi* (red dot), and human (black dot) indicated. Genus species and sequence identity for proteins used in the alignments are: *Crithidia fasciculata* (Gene ID CFAC1_060020300, CFAC1_300013300, CFAC1_270060100, CFAC1_290037400, CFAC1_290037500, CFAC1_290037600, CFAC1_290037700, CFAC1_290037800, CFAC1_290038200); human (UniProt PP1A_HUMAN, PP1B_HUMAN, PP1G_HUMAN); *Leishmania major* (Gene ID LmjF.15.0220, LmjF.28.0690, LmjF.31.2630, LmjF.34.0780, LmjF.34.0790, LmjF.34.0800, LmjF.34.0810, LmjF.34.0850); *Leishmania tartentolae* (Gene ID LtaP15.0230, LtaP28.0710, LtaP31.3050, LtaP34.0890, LtaP34.0920, LtaP34.0930, LtaP34.0940, LtaP34.0980); *Leptomonas seymouri* (Gene ID Lsey_0031_0170, Lsey_0328_0090, Lsey_0221_0060, Lsey_0565_0010 *Trypanosoma brucei* (Gene ID Tb927.4.3560, Tb927.4.3610, Tb927.4.3620, Tb927.4.3630, Tb927.4.3640, Tb927.4.5030, Tb927.8.7390, Tb927.11.8090); *Trypanosoma cruzi* (Gene ID TcCLB.506739.130, TcCLB.508815.110, TcCLB.506201.30, TcCLB.506201.70, TcCLB.506201.80, TcCLB.507757.50, TcCLB.509633.50); *Trypanosoma vivax* (Gene ID TvY486_0019980, TvY486_0403330, TvY486_0403390, TvY486_0806880, TvY486_1108810); *Trypanosoma grayi* (Gene ID DQ04_02191050, DQ04_11101010, DQ04_01081070, DQ04_02221000, DQ04_13331010, DQ04_16591000). The four *L. tarentolae* clade C genes are incomplete in the genome deposited at TriTrypDB, with each gene harboring internal ambiguous bases. Full-length copies of these genes derived from our in-house genome for *L. tarentolae* that were used in this figure. GeneIDs for these genes were assigned based on alignment with the incomplete copies available on TriTrypDB.

**FIG S3**

**Sequence analysis of identified proteins using HH****pred**. The schematics for each predicted protein show the length of the complete protein plus coordinates where the top hit of the protein database (PDB) aligned. Hits (predicted to be structurally similar to the query) are color-coded, with high probability hits in red and low probability hits in blue and black. Below the schematic showing the best hits are the HHpred descriptions of the corresponding proteins and similarities. This includes the PDB number and chain designation (Hit), a description of the PDB entry including related PDB entries (Name), the probability of the hit based on the Hidden Markov Model (Probability), the probability of the match in an unrelated database (E-value), score for the secondary structure prediction (SS), number of amino acids aligned (Col), and the total length of the target in PDB (Target Length). The documentation for HHpred considers “Probability” the most important criterion, with hits >50% being considered significant.

**FIG S4**

**Effect of depletion of JBP3 on transcription termination**.

The normalized read counts are shown for the 10 kb surrounding TTSs for T7-TR/JBP3-MHTAP/Δ*jbp3*::*neo* grown in the presence (JBP3+,blue line) or absence of Tet (JBP3-,orange line). Plots are oriented such that transcription is proceeds from the left and terminates at “0”, with the top strand being the coding strand on the left side of the TTS. The normalized read counts are shown. Panels on the left depict reads mapping to the top strand and panels on the right depict reads mapping to the bottom strand.

**A**. All TTSs.

**B**. TTSs in cSSRs lacking either an RNA gene or a centromere.

**C**. Unidirectional (“head to tail”) TTSs.

**D**. TTSs immediately upstream of an RNA gene.

**E**. TTSs adjacent to a centromere.

**F**. TTSs adjacent to a telomere.

**G**. Box-and-whiskers plots showing the median coverage in the 5 kb downstream of all TTSs (All), TTS at cSSRs (Conv), TTSs between head-to-tail PTUS (Uni), TTS immediately upstream of one or more RNA genes (RNA), TTS adjacent to a centromere (Cent) and TTSs at telomeres (Telo). Panel G (top strand) is also shown as FIG 5D.

**FIG S5**

**Effect of depletion of JBP3 on transcription initiation**.

A similar analysis as FIG S5, displaying the normalized read coverage for the 10 kb regions surrounding the transcription start sites. Plots are oriented such that transcription starts at position “0” and proceeds left to right, with the top strand being the coding strand to the right of the TSS. The normalized read counts for all TSSs mapping are shown. Panels on the left depict reads mapping to the top strand and panels on the right depict reads mapping to the bottom strand.

**A**. All TSSs.

**B**. TSSs in dSSRs lacking either an RNA gene or a centromere.

**C**. Unidirectional (“head to tail”) TSSs.

**D**. TSSs immediately downstream of an RNA gene.

**E**. TSSs adjacent to a centromere.

**F**. TSSs adjacent to a telomere.

**G**. Box-and-whiskers plots showing the median coverage in the 5 kb downstream of all TSSs (All), TSS at cSSRs (Div), TSSs between head-to-tail PTUS (Uni), TSSs immediately doownstream of one or more RNA genes (RNA), TSSs adjacent to a centromere (Cent) and TSSs at telomeres (Telo).

**FIG S6**

**Map of plasmid pLEXSY-MHTAP showing key features and restriction sites**.

Bait proteins were cloned into the listed restriction sites immediately upstream of the TAP tag.

**DATA SET S1**

Excel spreadsheet showing all proteins identified by mass spectrometry of TAP-tag pull-downs. The results from each experiment are shown in different tabs, with color shading used to indicate proteins that show substantial (>32-fold) enrichment over control. The final tab aggregates the results from all experiments with color shading used to differentiate proteins in separate complexes.

## References

1. S. Thomas, A. Green, N. R. Sturm, D. A. Campbell, P. J. Myler, Histone acetylations mark origins of polycistronic transcription in Leishmania major. BMC Genomics 10, 152 (2009).

2. S. Martinez-Calvillo et al., Transcription of Leishmania major Friedlin chromosome 1 initiates in both directions within a single region. Mol. Cell 11, 1291–1299 (2003).

3. S. Martinez-Calvillo, D. Nguyen, K. Stuart, P. J. Myler, Transcription initiation and termination on Leishmania major chromosome 3. Eukaryot Cell 3, 506–517 (2004).

4. J. H. Gommers-Ampt et al., β-D-glucosyl-hydroxymethyluracil: a novel modified base present in the DNA of the parasitic protozoan T. brucei. Cell 75, 1129–1136 (1993).

5. F. van Leeuwen, R. Kieft, M. Cross, P. Borst, Biosynthesis and function of the modified DNA base β-D-glucosyl-hydroxymethyluracil in Trypanosoma brucei. Mol. Cell. Biol. 18, 5643–5651 (1998).

6. F. van Leeuwen et al., β -D-glucosyl-hydroxymethyluracil is a conserved DNA modification in kinetoplastid protozoans and is abundant in their telomeres. Proc. Natl. Acad. Sci. U. S. A. 95, 2366–2371 (1998).

7. F. Van Leeuwen et al., The telomeric GGGTTA repeats of Trypanosoma brucei contain the hypermodified base J in both strands. Nucleic Acids Res. 24, 2476–2482 (1996).

8. F. van Leeuwen et al., Localization of the modified base J in telomeric VSG gene expression sites of Trypanosoma brucei. Genes Dev. 11, 3232–3241 (1997).

9. F. van Leeuwen, R. Kieft, M. Cross, P. Borst, Tandemly repeated DNA is a target for the partial replacement of thymine by β -D-glucosyl-hydroxymethyluracil in Trypanosoma brucei. Mol. Biochem. Parasitol. 109, 133–145 (2000).

10. L. J. Cliffe, T. N. Siegel, M. Marshall, G. A. Cross, R. Sabatini, Two thymidine hydroxylases differentially regulate the formation of glucosylated DNA at regions flanking polymerase II polycistronic transcription units throughout the genome of Trypanosoma brucei Nucleic Acids Res. 38, 3923–3935 (2010).

11. H. G. van Luenen et al., Glucosylated hydroxymethyluracil, DNA base J, prevents transcriptional readthrough in Leishmania. Cell 150, 909–921 (2012).

12. C. A. Marques, N. J. Dickens, D. Paape, S. J. Campbell, R. McCulloch, Genome-wide mapping reveals single-origin chromosome replication in Leishmania, a eukaryotic microbe. Genome Biol 16, 230 (2015).

13. M. R. Garcia-Silva et al., Identification of the centromeres of Leishmania major: revealing the hidden pieces. EMBO Rep 18, 1968–1977 (2017).

14. A. Sekar, C. Merritt, L. Baugh, K. Stuart, P. J. Myler, Tb927.10.6900 encodes the glucosyltransferase involved in synthesis of base J in Trypanosoma brucei. Mol. Biochem. Parasitol. 196, 9–11 (2014).

15. T. Heidebrecht et al., The structural basis for recognition of base J containing DNA by a novel DNA binding domain in JBP1. Nucleic Acids Res 39, 5715–5728 (2011).

16. C. DiPaolo, R. Kieft, M. Cross, R. Sabatini, Regulation of trypanosome DNA glycosylation by a SWI2/SNF2-like protein. Mol. Cell 17, 441–451 (2005).

17. L. J. Cliffe et al., JBP1 and JBP2 are two distinct thymidine hydroxylases involved in J biosynthesis in genomic DNA of African trypanosomes. Nucleic Acids Res 37, 1452–1462 (2009).

18. P. A. Genest et al., Formation of linear inverted repeat amplicons following targeting of an essential gene in Leishmania. Nucleic Acids Res 33, 1699–1709 (2005).

19. P. A. Genest et al., Defining the sequence requirements for the positioning of base J in DNA using SMRT sequencing. Nucleic Acids Res 43, 2102–2115 (2015).

20. R. Kieft et al., Identification of a novel base J binding protein complex involved in RNA polymerase II transcription termination in trypanosomes. PLoS Genet 16, e1008390 (2020).

21. M. Bollen, W. Peti, M. J. Ragusa, M. Beullens, The extended PP1 toolkit: designed to create specificity. Trends Biochem. Sci. 35, 450–458 (2010).

22. B. Lesage et al., A complex of catalytically inactive protein phosphatase-1 sandwiched between Sds22 and inhibitor-3. Biochemistry (Mosc). 46, 8909–8919 (2007).

23. L. Zimmermann et al., A Completely Reimplemented MPI Bioinformatics Toolkit with a New HHpred Server at its Core. J. Mol. Biol. 430, 2237–2243 (2018).

24. Y. M. Kim et al., PNUTS, a protein phosphatase 1 (PP1) nuclear targeting subunit. Characterization of its PP1-and RNA-binding domains and regulation by phosphorylation. J. Biol. Chem. 278, 13819–13828 (2003).

25. J. P. Kreivi et al., Purification and characterisation of p99, a nuclear modulator of protein phosphatase 1 activity. FEBS Lett. 420, 57–62 (1997).

26. P. B. Allen, Y. G. Kwon, A. C. Nairn, P. Greengard, Isolation and characterization of PNUTS, a putative protein phosphatase 1 nuclear targeting subunit. J. Biol. Chem. 273, 4089–4095 (1998).

27. J. H. Lee, J. You, E. Dobrota, D. G. Skalnik, Identification and characterization of a novel human PP1 phosphatase complex. J. Biol. Chem. 285, 24466–24476 (2010).

28. M. S. Choy et al., Understanding the antagonism of retinoblastoma protein dephosphorylation by PNUTS provides insights into the PP1 regulatory code. Proc. Natl. Acad. Sci. U. S. A. 111, 4097–4102 (2014).

29. H. B. Landsverk, M. Kirkhus, M. Bollen, T. Kuntziger, P. Collas, PNUTS enhances in vitro chromosome decondensation in a PP1-dependent manner. Biochem. J. 390, 709–717 (2005).

30. M. A. Cortazar et al., Control of RNA Pol II Speed by PNUTS-PP1 and Spt5 Dephosphorylation Facilitates Termination by a “Sitting Duck Torpedo” Mechanism. Mol. Cell 76, 896–908 e894 (2019).

31. S. C. Dillon, X. Zhang, R. C. Trievel, X. Cheng, The SET-domain protein superfamily: protein lysine methyltransferases. Genome Biol 6, 227 (2005).

32. R. Paro, Imprinting a determined state into the chromatin of Drosophila. Trends Genet. 6, 416–421 (1990).

33. R. Paro, D. S. Hogness, The Polycomb protein shares a homologous domain with a heterochromatin-associated protein of Drosophila. Proc. Natl. Acad. Sci. U. S. A. 88, 263–267 (1991).

34. R. Aasland, A. F. Stewart, The chromo shadow domain, a second chromo domain in heterochromatin-binding protein 1, HP1. Nucleic Acids Res 23, 3168–3173 (1995).

35. O. Rozenblatt-Rosen et al., The parafibromin tumor suppressor protein is part of a human Paf1 complex. Mol. Cell. Biol. 25, 612–620 (2005).

36. R. Pavri et al., Histone H2B monoubiquitination functions cooperatively with FACT to regulate elongation by RNA polymerase II. Cell 125, 703–717 (2006).

37. N. J. Krogan et al., RNA polymerase II elongation factors of Saccharomyces cerevisiae: a targeted proteomics approach. Mol. Cell. Biol. 22, 6979–6992 (2002).

38. B. A. Ouna et al., Depletion of trypanosome CTR9 leads to gene expression defects. PLoS One 7, e34256 (2012).

39. D. C. Scott et al., Blocking an N-terminal acetylation-dependent protein interaction inhibits an E3 ligase. Nat Chem Biol 13, 850–857 (2017).

40. L. C. Soares Medeiros et al., Rapid, selection-free, high-efficiency genome editing in protozoan parasites using CRISPR-Cas9 ribonucleoproteins. MBio 8, (2017).

41. H. Kim et al., TRF2 functions as a protein hub and regulates telomere maintenance by recognizing specific peptide motifs. Nat Struct Mol Biol 16, 372–379 (2009).

42. A. Ciurciu et al., PNUTS/PP1 regulates RNAPII-mediated gene expression and is necessary for developmental growth. PLOS Genetics 9, e1003885 (2013).

43. C. U. Stirnimann, E. Petsalaki, R. B. Russell, C. W. Muller, WD40 proteins propel cellular networks. Trends Biochem. Sci. 35, 565–574 (2010).

44. M. Schapira, M. Tyers, M. Torrent, C. H. Arrowsmith, WD40 repeat domain proteins: a novel target class? Nat Rev Drug Discov 16, 773–786 (2017).

45. S. Dean, J. D. Sunter, R. J. Wheeler, TrypTag.org: A Trypanosome genome-wide protein localisation resource. Trends Parasitol 33, 80–82 (2017).

46. A. Yart et al., The HRPT2 tumor suppressor gene product parafibromin associates with human PAF1 and RNA polymerase II. Mol. Cell. Biol. 25, 5052–5060 (2005).

47. B. N. Tomson, K. M. Arndt, The many roles of the conserved eukaryotic Paf1 complex in regulating transcription, histone modifications, and disease states. Biochim Biophys Acta 1829, 116–126 (2013).

48. S. B. Van Oss, C. E. Cucinotta, K. M. Arndt, Emerging insights into the roles of the Paf1 Complex in gene regulation. Trends Biochem. Sci. 42, 788–798 (2017).

49. C. L. Mueller, J. A. Jaehning, Ctr9, Rtf1, and Leo1 are components of the Paf1/RNA polymerase II complex. Mol. Cell. Biol. 22, 1971–1980 (2002).

50. B. Zhu et al., The human PAF complex coordinates transcription with events downstream of RNA synthesis. Genes Dev. 19, 1668–1673 (2005).

51. M. K. Mayekar, R. G. Gardner, K. M. Arndt, The recruitment of the Saccharomyces cerevisiae Paf1 complex to active genes requires a domain of Rtf1 that directly interacts with the Spt4-Spt5 complex. Mol. Cell. Biol. 33, 3259–3273 (2013).

52. Y. Xie et al., Paf1 and Ctr9 subcomplex formation is essential for Paf1 complex assembly and functional regulation. Nat Commun 9, 3795 (2018).

53. T. Kurz et al., The conserved protein DCN-1/Dcn1p is required for cullin neddylation in C. elegans and S. cerevisiae. Nature 435, 1257–1261 (2005).

54. Y. Nakazawa et al., Ubiquitination of DNA damage-stalled RNAPII promotes transcription-coupled repair. Cell 180, 1228–1244 e1224 (2020).

55. A. Tufegdzic Vidakovic et al., Regulation of the RNAPII pool is integral to the DNA damage response. Cell 180, 1245–1261 e1221 (2020).

56. B. C. Jensen et al., Characterization of protein kinase CK2 from Trypanosoma brucei. Mol. Biochem. Parasitol. 151, 28–40 (2007).

57. G. M. Kapler, K. Zhang, S. M. Beverley, Nuclease mapping and DNA sequence analysis of transcripts from the dihydrofolate reductase-thymidylate synthase (R) region of Leishmania major Nucleic Acids Res. 18, 6399–6408 (1990).

58. H. L. Parker et al., Three genes and two isozymes: Gene conversion and the compartmentalization and expression of the phosphoglycerate kinases of Trypanosoma (Nannomonas) congolense Mol. Biochem. Parasitol. 69, 269–279 (1995).

59. A. Michalski et al., Ultra high resolution linear ion trap Orbitrap mass spectrometer (Orbitrap Elite) facilitates top down LC MS/MS and versatile peptide fragmentation modes. Molecular & cellular proteomics : MCP 11, O111 013698 (2012).

60. J. Cox, M. Mann, MaxQuant enables high peptide identification rates, individualized p.p.b.-range mass accuracies and proteome-wide protein quantification. Nat. Biotechnol. 26, 1367–1372 (2008).

61. J. Cox et al., Accurate proteome-wide label-free quantification by delayed normalization and maximal peptide ratio extraction, termed MaxLFQ. Molecular & cellular proteomics : MCP 13, 2513–2526 (2014).

62. S. Tyanova et al., The Perseus computational platform for comprehensive analysis of (prote)omics data. Nat Methods 13, 731–740 (2016).

63. P. Di Tommaso et al., T-Coffee: a web server for the multiple sequence alignment of protein and RNA sequences using structural information and homology extension. Nucleic Acids Res 39, W13–17 (2011).

64. Y. Song et al., High-resolution comparative modeling with RosettaCM. Structure 21, 1735–1742 (2013).

65. T. Beneke et al., A CRISPR Cas9 high-throughput genome editing toolkit for kinetoplastids. R Soc Open Sci 4, 170095 (2017).

66. G. Schumann Burkard, P. Jutzi, I. Roditi, Genome-wide RNAi screens in bloodstream form trypanosomes identify drug transporters. Mol. Biochem. Parasitol. 175, 91–94 (2011).

67. A. G. Hunt, A rapid, simple, and inexpensive method for the preparation of strand-specific RNA-Seq libraries. Methods Mol. Biol. 1255, 195–207 (2015).

68. P. J. Myler, J. A. McDonald, P. J. Alcolea, A. Sur, Quantitative RNA analysis using RNA-Seq. Methods Mol. Biol. 1971, 95–108 (2019).

69. B. Langmead, S. L. Salzberg, Fast gapped-read alignment with Bowtie 2. Nat Methods 9, 357–359 (2012).

