## Supplementary Table S1 for "Chromatin-associated protein complexes link DNA base J and transcription termination in *Leishmania*"

### Table S1. Protein complexes identified by TAP-tagged experiments

| Protein name | | MW | TriTrypDB annotation (Gene ID) | # of peptides | | $log_2$ fold-enrichment | | |
|---|---|---|---|---|---|---|---|---|
| Bait | Pull-down | (kDa) | | Rep 1 | Rep 2 | Rep 1 | Rep 2 | Avg |
| HmdUGT | HmdUGT | 101.3 | Hypothetical protein, conserved (LtaP36.2450) | 55 | 41 | 13.7 | 14.6 | 14.1 |
| | JBP3 | 73.9 | Hypothetical protein, conserved (LtaP36.0380) | 19 | 7 | 13.6 | 8.6 | 11.1 |
| | WD-GT | 38.6 | Hypothetical protein, conserved (LtaP32.3990) | 13 | 7 | 11.7 | 6.9 | 9.3 |
| | PP1Ce | 42.3 | Protein phosphatase 1 catalytic subunit, putative (LtaP15.0230) | 10 | 3 | 9.8 | 5.1 | 7.4 |
| | PNUTS | 28.6 | Hypothetical protein, conserved (LtaP33.1440) | 8 | 0 | 9.8 | - | 4.9 |
| | PFDN3 | 22.3 | Prefoldin-like protein (LtaP26.1310) | 6 | 4 | 8.8 | 7.8 | 8.3 |
| | PFDN6 | 15.8 | Prefoldin subunit, putative (LtaP05.1280) | 4 | 3 | 8.0 | 6.0 | 7.0 |
| | PFDN5 | 18.0 | Prefoldin 5-like protein (LtaP22.0650) | 3 | 2 | 8.4 | 5.5 | 6.9 |
| | PFDN4 | 14.9 | Hypothetical protein, conserved (LtaP25.0870) | 2 | 2 | 7.0 | 5.7 | 6.4 |
| | PFDN1 | 16.5 | Hypothetical protein, conserved (LtaP30.0960) | 4 | 2 | 6.3 | 3.7 | 5.0 |
| | PFDN2 | 15.3 | Prefoldin subunit 2, putative (LtaP09.0680) | 1 | 0 | 5.6 | - | 2.8 |
| PP1Ce | PP1Ce | 42.3 | Protein phosphatase 1 catalytic subunit, putative (LtaP15.0230) | 16 | 21 | 17.1 | 17.1 | 17.1 |
| | WD-GT | 38.6 | Hypothetical protein, conserved (LtaP32.3990) | 21 | 25 | 12.6 | 17.1 | 14.8 |
| | PNUTS | 28.6 | Hypothetical protein, conserved (LtaP33.1440) | 15 | 15 | 12.9 | 12.1 | 12.5 |
| | JBP3 | 73.9 | Hypothetical protein, conserved (LtaP36.0380) | 28 | 34 | 11.8 | 12.8 | 12.3 |
| | HmdUGT | 101.3 | Hypothetical protein, conserved (LtaP36.2450) | 23 | 25 | 4.8 | 9.6 | 7.2 |
| | PPP1R7 | 44.7 | Protein phosphatase type 1 regulator-like protein (LtaP05.1290) | 26 | 35 | 15.8 | 15.0 | 15.4 |
| | PPP1R11 | 16.5 | Protein phosphatase inhibitor, putative (LtaP07.0770) | 11 | 7 | 14.3 | 13.6 | 13.9 |
| | PPP1R2 | 15.5 | Protein phosphatase inhibitor 2, IPP-2, putative (LtaP29.0170) | 8 | 8 | 11.7 | 12.9 | 12.3 |
| | DNAJ | 43.7 | Heat shock protein DNAJ, putative (LtaP27.2520) | 6 | 9 | 7.7 | 5.6 | 6.6 |
| PNUTS | PNUTS | 28.6 | Hypothetical protein, conserved (LtaP33.1440) | 16 | 17 | 17.1 | 18.2 | 17.7 |
| | WD-GT | 38.6 | Hypothetical protein, conserved (LtaP32.3990) | 22 | 22 | 12.2 | 13.2 | 12.7 |
| | PP1Ce | 42.3 | Protein phosphatase 1 catalytic subunit, putative (LtaP15.0230) | 16 | 15 | 12.1 | 12.3 | 12.2 |
| | JBP3 | 73.9 | Hypothetical protein, conserved (LtaP36.0380) | 30 | 24 | 10.0 | 10.6 | 10.3 |
| | HmdUGT | 101.3 | Hypothetical protein, conserved (LtaP36.2450) | 14 | 14 | 9.4 | 7.0 | 8.2 |
| WD-GT | WD-GT | 38.6 | Hypothetical protein, conserved (LtaP32.3990) | 21 | 25 | 13.9 | 18.6 | 16.3 |
| | PP1Ce | 42.3 | Protein phosphatase 1 catalytic subunit, putative (LtaP15.0230) | 15 | 17 | 13.3 | 14.6 | 13.9 |
| | PNUTS | 28.6 | Hypothetical protein, conserved (LtaP33.1440) | 14 | 15 | 13.5 | 13.0 | 13.2 |
| | JBP3 | 73.9 | Hypothetical protein, conserved (LtaP36.0380) | 32 | 36 | 11.6 | 13.5 | 12.5 |
| | HmdUGT | 101.3 | Hypothetical protein, conserved (LtaP36.2450) | 26 | 23 | 8.1 | 10.3 | 9.2 |
| | PFDN6 | 15.8 | Prefoldin subunit, putative (LtaP05.1280) | 8 | 10 | 10.8 | 9.5 | 10.2 |
| | PFDN2 | 15.3 | Prefoldin subunit 2, putative (LtaP09.0680) | 3 | 4 | 10.2 | 6.7 | 8.4 |
| | PFDN5 | 18.0 | Prefoldin 5-like protein (LtaP22.0650) | 3 | 5 | 9.4 | 8.4 | 8.9 |
| | PFDN4 | 14.9 | Prefoldin subunit, putative (LtaP25.0870) | 3 | 3 | 8.8 | 8.8 | 8.8 |
| | PFDN1 | 16.5 | Prefoldin subunit, putative (LtaP30.0960) | 7 | 3 | 8.8 | 5.5 | 7.1 |
| | PFDN3 | 22.3 | Prefoldin-like protein (LtaP26.1310) | 6 | 10 | 7.2 | 10.0 | 8.6 |
| | CCT7 | 61.7 | T-complex protein 1, eta subunit, putative (LtaP35.3870) | 24 | 28 | 12.0 | 5.8 | 8.9 |
| | CCT5 | 94.4 | Chaperonin containing t-complex protein, putative (LtaP32.1090) | 19 | 18 | 11.2 | 4.9 | 8.1 |
| | CCT8 | 58.3 | TCP-1/cpn60 chaperonin family, putative (LtaP36.7170) | 26 | 18 | 10.3 | 7.7 | 9.0 |
| | CCT3 | 60.1 | T-complex protein 1, gamma subunit, putative (LtaP23.1490) | 20 | 21 | 10.1 | 8.3 | 9.2 |
| | CCT1 | 59.2 | Chaperonin alpha subunit, putative (LtaP32.3480) | 9 | 11 | 7.3 | 6.7 | 7.0 |
| | CCT4 | 59.6 | T-complex protein 1, delta subunit, putative (LtaP21.1260) | 18 | 17 | 7.0 | 5.5 | 6.2 |
| | DNAJ | 43.7 | Heat shock protein DNAJ, putative (LtaP27.2520) | 8 | 9 | 9.0 | 5.6 | 7.3 |
| | HSP110 | 91.1 | Heat shock protein 110, putative (LtaP18.1330) | 8 | 15 | 6.8 | 7.5 | 7.1 |
| | ZFK | 144.0 | Zinc finger protein kinase-like (LtaP28.1730) | 10 | 6 | 8.0 | 5.4 | 6.7 |
| | - | 161.3 | Hypothetical protein, conserved (LtaP33.2370) | 14 | 12 | 7.7 | 6.1 | 6.9 |
| JBP3 | JBP3 | 73.9 | Hypothetical protein, conserved (LtaP36.0380) | 38 | 44 | 14.9 | 17.0 | 15.9 |
| | SET-J3C | 88.8 | SET domain containing protein, putative (LtaP35.2400) | 18 | 22 | 12.6 | 13.0 | 12.8 |
| | CS-J3C | 24.1 | Hypothetical protein, conserved (LtaP28.2640) | 10 | 8 | 13.5 | 11.5 | 12.5 |
| | HPC-J3C[c] | 93.5 | Hypothetical protein, conserved (LtaP12.0900) | 26 | 31 | 11.8 | 13.1 | 12.4 |
| | Chromo-J3C | 40.2 | Chromo domain-containing protein (LtaP14.0150) | 12 | 18 | 12.6 | 10.8 | 11.7 |
| | WD-GT | 38.6 | Hypothetical protein, conserved (LtaP32.3990) | 14 | 24 | 8.0 | 16.6 | 12.3 |
| | PP1Ce | 42.3 | Protein phosphatase 1 catalytic subunit, putative (LtaP15.0230) | 13 | 16 | 8.8 | 12.9 | 10.9 |
| | PNUTS | 28.6 | Hypothetical protein, conserved (LtaP33.1440) | 11 | 15 | 9.0 | 11.6 | 10.3 |
| | HmdUGT | 101.3 | Hypothetical protein, conserved (LtaP36.2450) | 30 | 27 | 9.6 | 11.6 | 10.6 |
| | LEO1 | 63.2 | RNA polymerase-associated protein, putative (LtaP35.2870) | 11 | 12 | 8.1 | 6.8 | 7.4 |
| | DCNL | 68.6 | Hypothetical protein, conserved (LtaP29.1270) | 6 | 2 | 6.4 | 4.5 | 5.4 |
| | CTR9 | 97.1 | RNA polymerase-associated protein CTR9, putative (LtaP29.2750) | 2 | 0 | 6.2 | - | 3.1 |
| | CDC73 | 41.0 | RNA polymerase-associated protein CDC73, putative (LtaP36.4090) | 5 | 1 | 5.7 | -0.5 | 2.6 |
| Chromo-J3C | Chromo-J3C | 40.2 | Chromo domain containing protein (LtaP14.0150) | 26 | 26 | 11.6 | 11.6 | 11.6 |
| | HPC-J3C[c] | 93.5 | Hypothetical protein, conserved (LtaP12.0900) | 26 | 26 | 10.4 | 9.2 | 9.8 |
| | CS-J3C | 24.1 | Hypothetical protein, conserved (LtaP28.2640) | 11 | 13 | 10.0 | 9.5 | 9.8 |
| | SET-J3C | 88.8 | SET domain containing protein, putative (LtaP35.2400) | 20 | 19 | 8.8 | 7.5 | 8.2 |
| | JBP3 | 73.9 | Hypothetical protein, conserved (LtaP36.0380) | 26 | 25 | 8.6 | 7.1 | 7.9 |
| CS-J3C | CS-J3C | 24.1 | Hypothetical protein, conserved (LtaP28.2640) | 11 | 13 | 11.8 | 11.9 | 11.8 |
| | SET-J3C | 88.8 | SET domain containing protein, putative (LtaP35.2400) | 21 | 20 | 9.6 | 9.3 | 9.4 |
| | Chromo-J3C | 40.2 | Chromo domain containing protein (LtaP14.0150) | 23 | 21 | 9.1 | 8.5 | 8.8 |
| | JBP3 | 73.9 | Hypothetical protein, conserved (LtaP36.0380) | 20 | 17 | 8.0 | 7.5 | 7.8 |
| | HPC-J3C[c] | 93.5 | Hypothetical protein, conserved (LtaP12.0900) | 29 | 27 | 6.2 | 6.4 | 6.3 |
| | - | 21.1 | i/6 autoantigen-like protein (LtaP22.1440) | 4 | 6 | 4.6 | 7.8 | 6.2 |
| LEO1 | LEO1 | 63.2 | RNA polymerase-associated protein, putative (LtaP35.2870) | 33 | 34 | 17.0 | 16.6 | 16.8 |
| | DCNL | 68.6 | Hypothetical protein, conserved (LtaP29.1270) | 35 | 35 | 14.0 | 13.8 | 13.9 |
| | CDC73 | 41.0 | RNA polymerase-associated protein (CDC73), putative (LtaP36.4090) | 11 | 13 | 10.3 | 10.4 | 10.4 |
| | CTR9 | 97.1 | RNA polymerase-associated protein (CTR9), putative (LtaP29.2750) | 28 | 30 | 9.6 | 9.3 | 9.5 |
| | RTF1L | 73.5 | Hypothetical protein, conserved (LtaP14.0860) | 18 | 19 | 4.2 | 3.5 | 3.8 |
| | PEX12 | 50.8 | Hypothetical protein, conserved (LtaP13.1250) | 5 | 5 | 6.2 | 4.6 | 5.4 |
| | TFIIS2-1 | 50.9 | Transcription elongation factor-like protein (LtaP33.3050) | 4 | 6 | 5.2 | 5.1 | 5.1 |
