## Supplementary Table S3 for "Chromatin-associated protein complexes link DNA base J and transcription termination in *Leishmania*"

### Table S3. Oligonucleotide primers used for construct creation.

| Primer | Sequence (5'-3') |
|---|---|
| **Primers used for creation of pLEXY-MHTAP** | |
| MHTAP-BamHI-S | GGATCCCCTAGGGGTACCCAATTGCTCGAGATGGAACAGAAACTGATCTCTG |
| MHTAP-NotI-AS | GCGGCCGCATGGGCAGGATCAGGTTGAC |
| **Primers used to generate TAP-tagged proteins** | |
| HmdUGT-AvrII-ATG | CGGCCTAGGATGTTCTCCCTCAACATCAAG |
| HmdUGT-SalI-P2790M | CGCGTCGACTGGACCAGGCTCGCCCTC |
| WD-GT-AvrII-ATG | CGGCCTAGGATGAACAGCTCCCCCGCCCCAAG |
| WD-GT-SalI-P1161M | CGCGTCGACTGTCTGTCCCTCCTTCAGTGATG |
| JBP3-AvrII-M9P | CGGCCTAGGCTTCTCAGCATGTCCTCAAAAC |
| JBP3-XhoI-P1989M | CGGCTCGAGTGTTTGTGAGGACGCCAGGCTC |
| PP1Ce-AvrII-ATG | CGGCCTAGGATGGCAAGCACGAAAAGAG |
| PP1Ce-MunI-P1122M | CGGCAATTGGTCGTGGCTCAAAGGATTTG |
| PNUTS-AvrII-ATG | GCACCTAGGCCATGAGCGTGGAGGAACTG |
| PNUTS-XhoI-P796M | GCACTCGAGCATGGGGATGGGGAGGGGCTC |
| LEO1-AvrII-ATG | GCACCTAGGTGGAATGGAGTGCCAAGCAG |
| LEO1-SalI-P1701M | GCAGTCGACCAGCTCACCAGGAAACAGTG |
| Chromo-J3C-AvrII-ATG | GCACCTAGGATGACCTACTACACCGTTGAG |
| Chromo-J3C-XhoI-P1080M | GCACTCGAGATGCAGAACGACCGAGTTC |
| CS-J3C-AvrII-ATG | GCACCTAGGATGCACGACGATCAGCTGCGCTTC |
| CS-J3-SalI-P684M | GCAGTCGACCGCCCAAACAGACATCGAGAG |
| **Primers used for CRISPR/Cas9 deletions** | |
| JBP3-up | **TGTGCACTACCCACCGTCTTTAGAACTCGG**GTATAATGCAGACCTGCTGC |
| JBP3-down | **CTTCCTCCGTGGCCCTCATGTGCGACTCCT**CCAATTTGAGAGACCTGTGC |
| JBP3-M84P | ACCCACCGTCTTTAGAACTC |
| JBP3-P2147M | TCTCTGATGGGCGTACATTC |
