## Supplementary figures and images for "Chromatin-associated protein complexes link DNA base J and transcription termination in *Leishmania*"

### Supplementary Figure S1

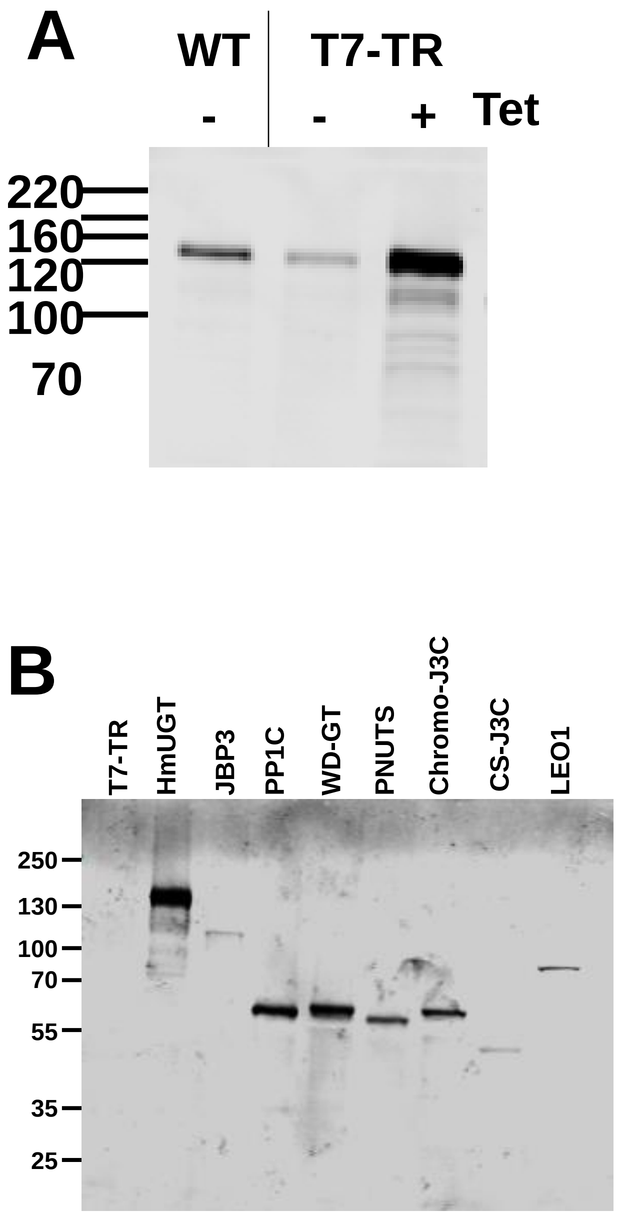

### Supplementary Figure S4

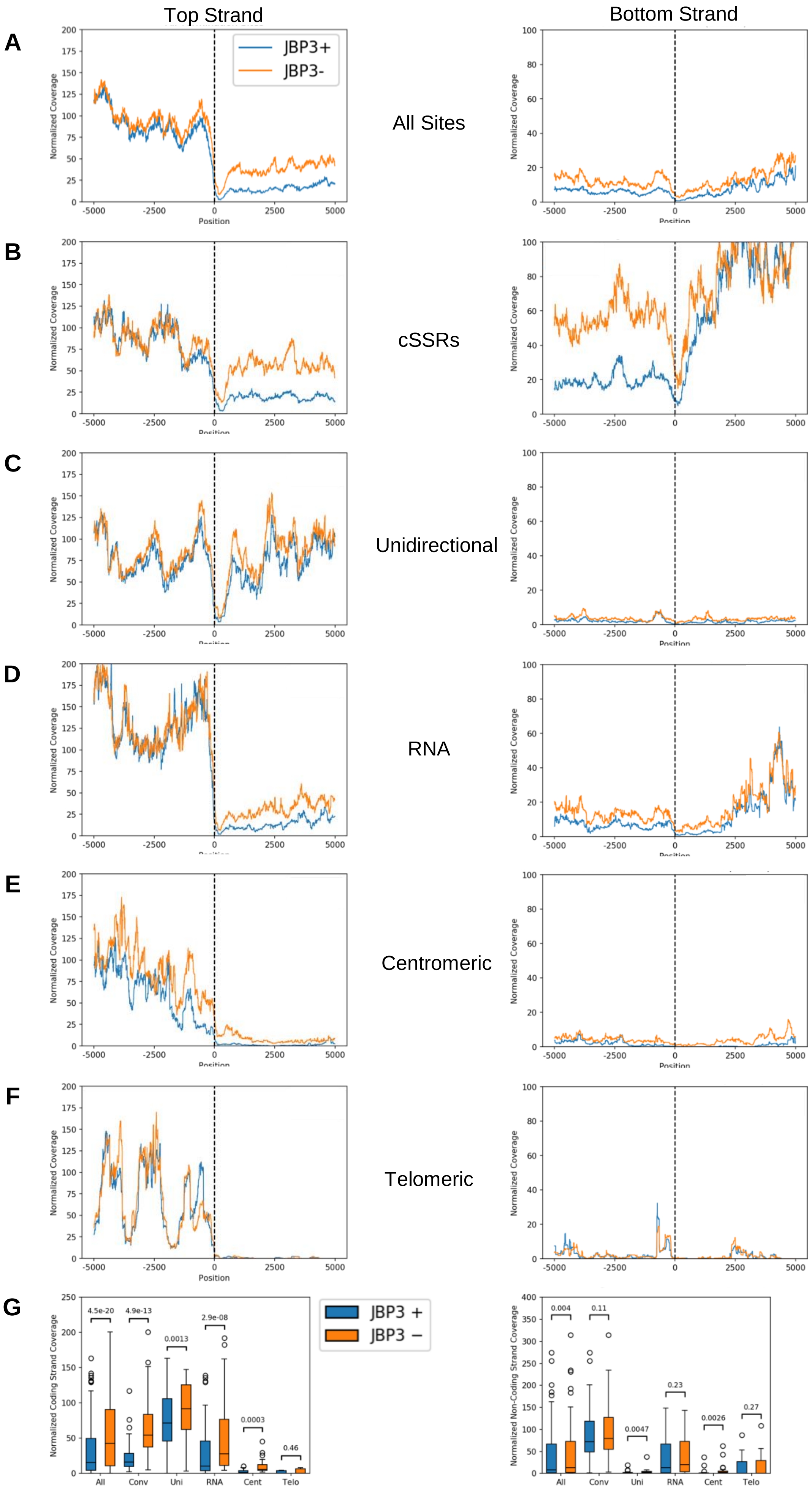

### Supplementary Figure S5

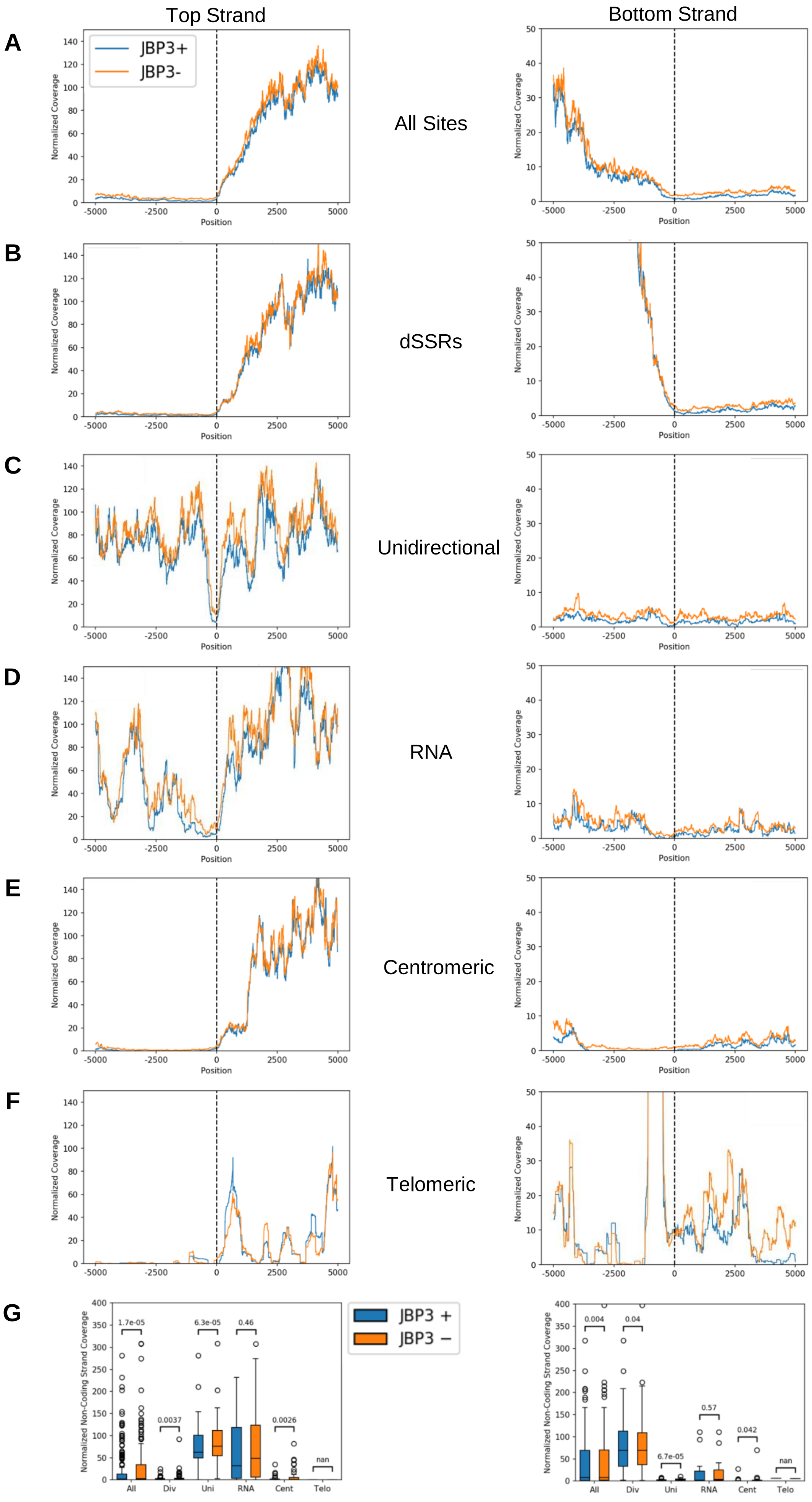

### Supplementary Figure S6

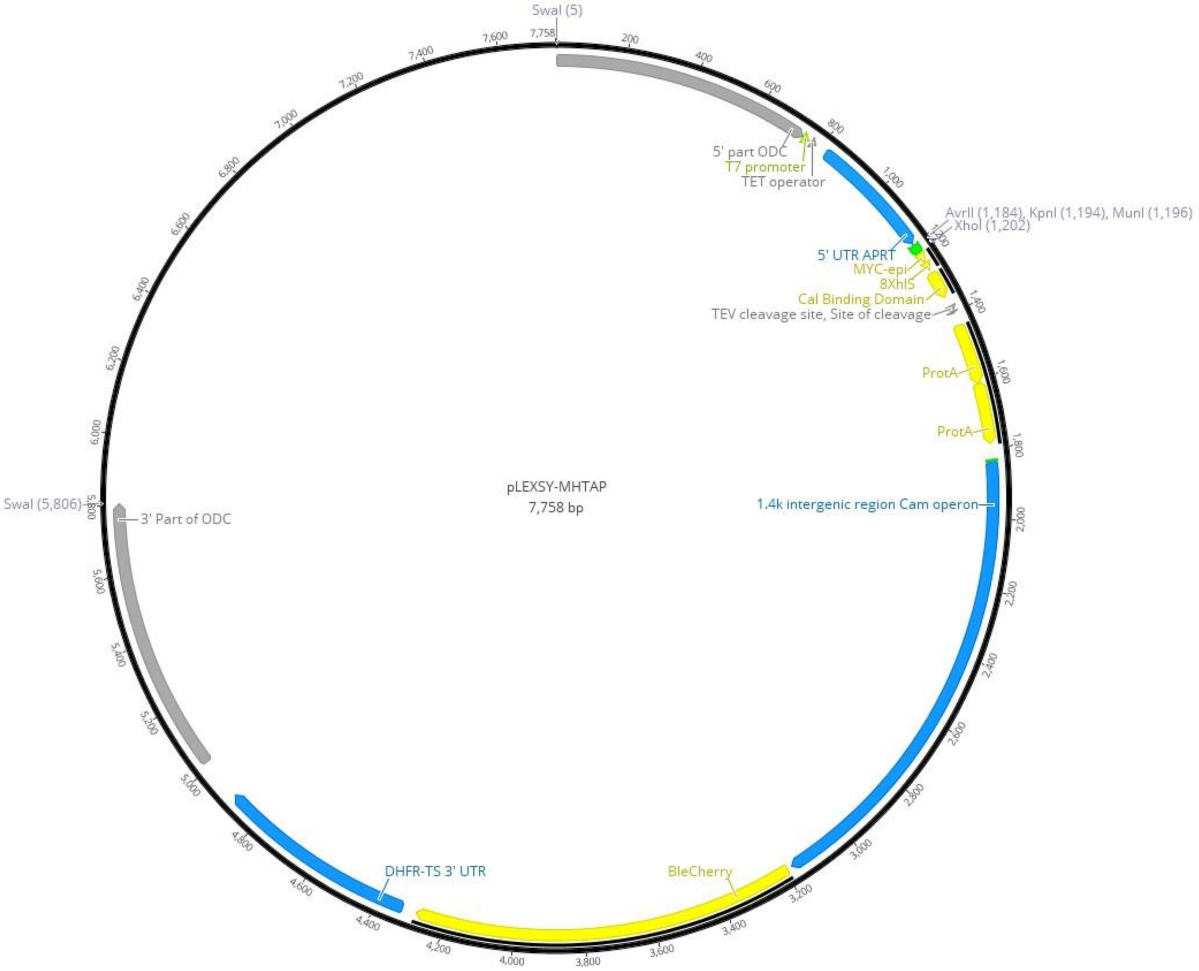
