## Supplementary Figure S2 for "Chromatin-associated protein complexes link DNA base J and transcription termination in *Leishmania*"

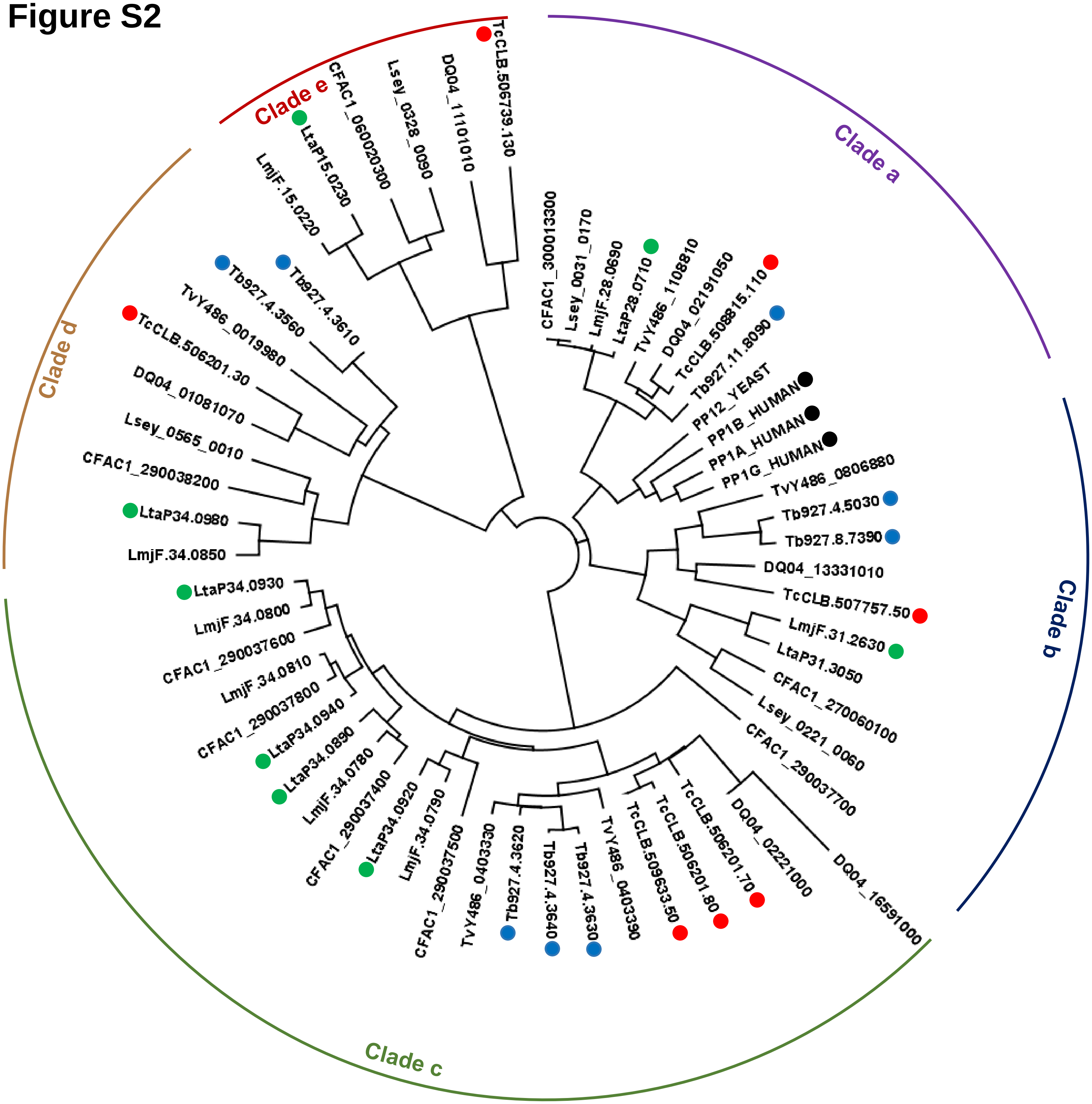


| | |
|---|---|
| Leishmaniiae | *Crithidia fasiculata* (CFAC)<br>*Leishmania major* (LmjF)<br>*Leptomonas seymouri* (Lsey)<br>*Leishmani tarentolae* (LtaP) ● (green) |
| Stercorarian trypanosomes | *Trypanosoma cruzi* (TcCLB) ● (red)<br>*Trypanosoma grayi* (DQ04) |
| Salivarian trypanosomes | *Trypanosoma brucei* (Tb927) ● (blue)<br>*Trypanosoma vivax* (TvY) |
| | *Homo sapiens* (Human) ● (black)<br>*Saccharomyces cerevisiae* (Yeast) |
