## Supplementary Figure S3 for "Chromatin-associated protein complexes link DNA base J and transcription termination in *Leishmania*"

### Figure S3A

#### PNUTS

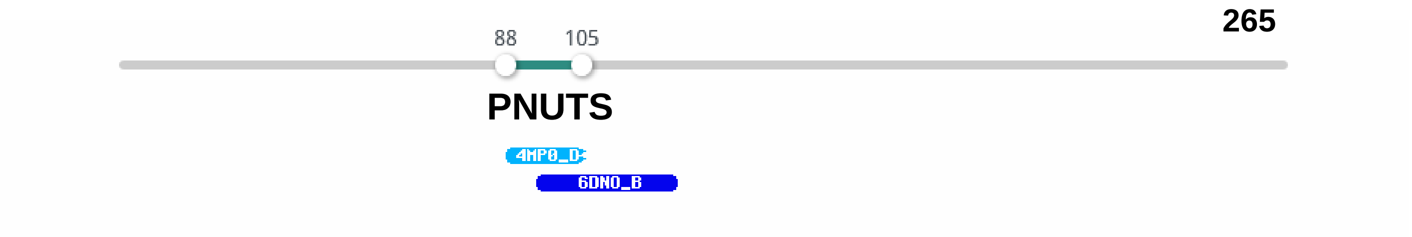


#### PNUTS

| Hit | Name | Probability | E-value | SS | Cols | Target Length |
|---|---|---|---|---|---|---|
| 4MP0_D | Serine/threonine-protein phosphatase PP1-alpha catalytic subunit; Serine/threonine phosphatase, Nucleus, HYDROLASE; HET: GOL, PO4; 2.1003A {Homo sapiens}; Related PDB entries: 4MOY_B 4MP0_B | 55.07 | 9.2 | 1.6 | 18 | 44 |
| 6DNO_B | Serine/threonine-protein phosphatase PP1-alpha catalytic subunit; Inhibitor, complex, HYDROLASE, HYDROLASE-HYDROLASE INHIBITOR; HET: 1ZN; 1.45A {Homo sapiens} | 38.83 | 30 | 2 | 30 | 30 |

### Figure S3B

## WD-GT

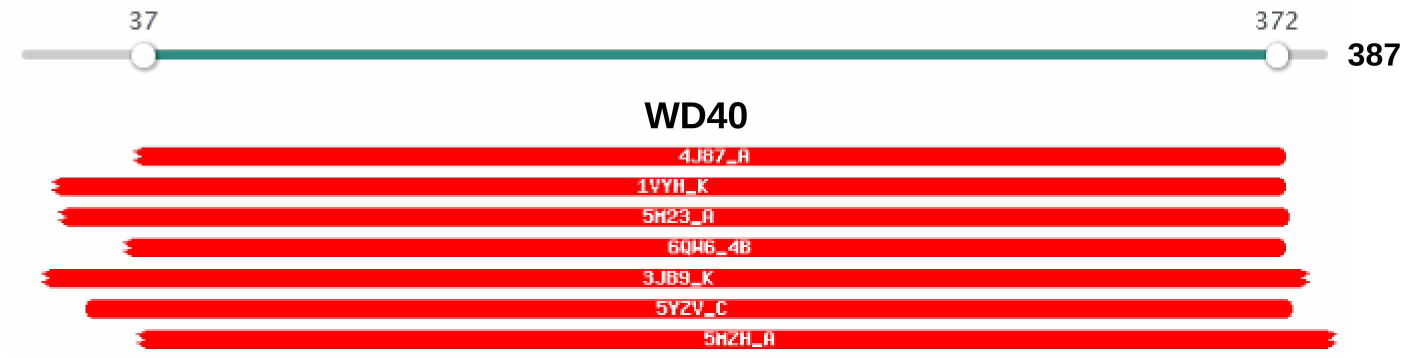


## WD40

| Hit | Name | Probability | E-value | SS | Cols | Target Length |
|---|---|---|---|---|---|---|
| 4J87_A | coatomer subunit alpha; Beta propeller domain, Vesicle trafficking; 1.67A {Schizosaccharomyces pombe}; Related PDB entries: 6PBG_A 4J8G_B 4J8G_A 4J8B_A | 100 | 1.1e-33 | 36.5 | 285 | 327 |
| 1VYH_K | PLATELET-ACTIVATING FACTOR ACETYLHYDROLASE IB BETA; LISSENCEPHALY, ACETYLHYDROLASE, PLATELET ACTIVACTING FACTOR; 3.4A {HOMO SAPIENS} SCOP: b.69.4.1; Related PDB entries: 1VYH_D 1VYH_C 1VYH_T 1VYH_P 1VYH_L 1VYH_H 1VYH_S 1VYH_O 1VYH_G | 100 | 3.5e-33 | 40.5 | 307 | 410 |
| 5M23_A | WD repeat-containing protein 5; MLL1 Methyltransferase peptide complex WDR5; HET: 7DC; 1.97A {Homo sapiens} SCOP: b.69.4.0; Related PDB entries: 5M25_A | 100 | 1.8e-33 | 37.9 | 310 | 434 |
| 6QW6_4B | Small nuclear ribonucleoprotein Sm D1; RNP complex, splicing, RNA, protein; HET: M7M, GTP, IHP; 2.92A {Homo sapiens}; Related PDB entries: 6AH0_K 6QX9_4B 6AHD_K 5O9Z_F 3JCR_L | 100 | 6.3e-33 | 39.6 | 286 | 522 |
| 3JB9_K | Pre-mRNA-splicing factor spp42, Pre-mRNA-splicing factor; Spliceosome, U2/U5/U6, Lariat, RNA BINDING; HET: ADP, GDP; 3.6A {Schizosaccharomyces pombe 972h-} SCOP: b.69.4.1 | 100 | 2.6e-32 | 42.4 | 313 | 473 |
| 5YZV_C | Probable serine/threonine-protein kinase PkwA (E.C.2.7.11.1); prokaryotic, thermostable, reversible folding, PROTEIN; 2.6A {Thermomonospora curvata}; Related PDB entries: 5YZV_B 5YZV_E 5YZV_D 5YZV_A | 100 | 2e-32 | 39 | 299 | 302 |
| 5MZH_A | Dynein assembly factor with WDR; Intraflagellar transport, cilium, beta propeller; HET: SO4; 2.398A {Chlamydomonas reinhardtii}; Related PDB entries: 5MZH_B | 100 | 1.3e-32 | 39.8 | 296 | 452 |

### Figure S3C

#### JBP3

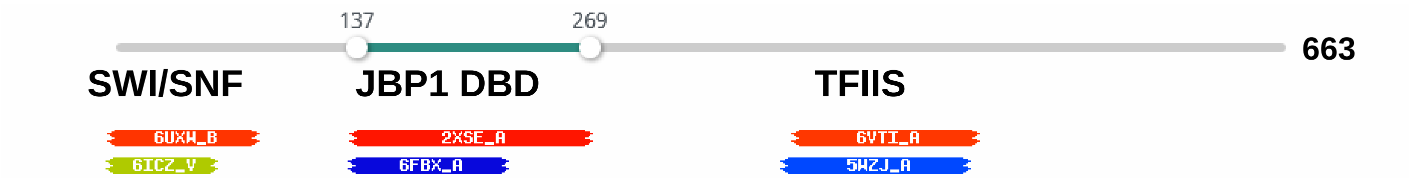


#### SWI/SNF

| | | | | | | |
|---|---|---|---|---|---|---|
| 6UXW_B | Histone H3.2, Histone H4, Histone; SWI/SNF, chromatin remodeler, TRANSCRIPTION, TRANSCRIPTION-DNA; HET: ADP, PO4;{Saccharomyces cerevisiae (strain ATCC 204508 / S288c)}; Related PDB entries: 6UXV_B | 95.6 | 0.011 | 4.1 | 82 | 1314 |
| 6ICZ_V | Protein mago nashi homolog 2; Human Post-catalytic Spliceosome, SPLICING; HET: SEP, ATP, GTP, I6P; 3.0A {Homo sapiens}; Related PDB entries: 6QDV_H 6FF7_T 5Z57_V 5XJC_V 5MQF_T 5Z56_V 5Z58_V 5YZG_V | 80.3 | 1.2 | 2.6 | 60 | 908 |

#### JBP1 DBD

| Hit | Name | Probability | E-value | SS | Cols | Target Length |
|---|---|---|---|---|---|---|
| 2XSE_A | THYMINE DIOXYGENASE JBP1 (E.C.1.14.11.6); OXIDOREDUCTASE, DNA-BINDING; HET: MSE; 1.9A {LEISHMANIA TARENTOLAE} | 98.2 | 0.000011 | 10.5 | 122 | 170 |
| 6FBX_A | BCL2-like 10, BCL2-antagonist of cell; Bcl-2 family, pro-survival protein, APOPTOSIS; 1.639A {Danio rerio}; Related PDB entries: 6H1N_A | 37.57 | 210 | 7.9 | 81 | 152 |

#### TFIIS

| | | | | | | |
|---|---|---|---|---|---|---|
| 6VTI_A | E3 ubiquitin-protein ligase UHRF1 (E.C.2.3.2.27); PP1-PNUTS phosphatase complex, Regulatory Subunit; NMR {Rattus norvegicus} | 95.26 | 0.13 | 8.7 | 98 | 148 |
| 5WZJ_A | Pumilio homolog 23, RNA; RNA binding protein, SIGNALING PROTEIN; 2.101A {Arabidopsis thaliana}; Related PDB entries: 5WZK_A 5WZH_A 5WZI_A 5WZG_A | 46.13 | 180 | 10.3 | 96 | 582 |

### Figure S3D

#### SET-J3C

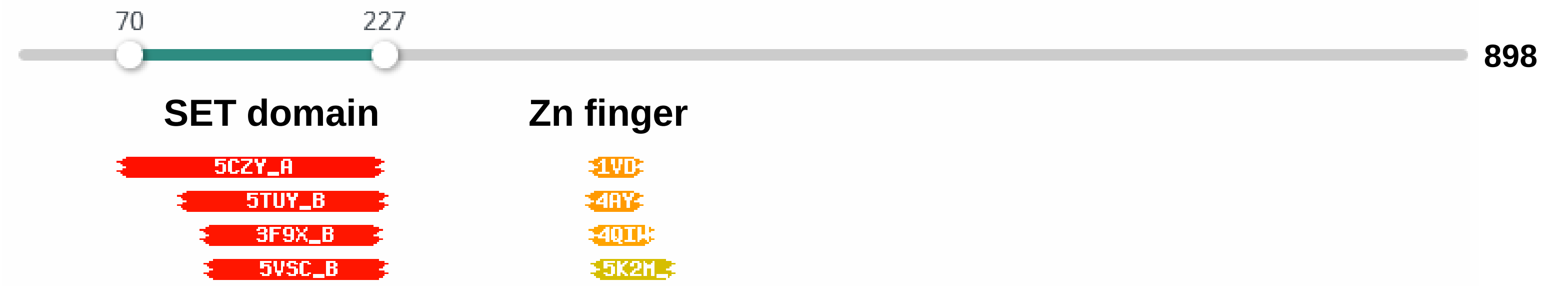


#### SET domain

| Hit | Name | Probability | E-value | SS | Cols | Length |
|---|---|---|---|---|---|---|
| 5CZY_A | Eukaryotic huntingtin interacting protein B; SET domain, TRANSFERASE; HET: SAM, GOL; 2.2A {Legionella pneumophila subsp. pneumophila str. Philadelphia 1} | 98.55 | 1.8e-8 | 0.8 | 134 | 483 |
| 5TUY_B | Histone-lysine N-methyltransferase EHMT2 (E.C.2.1.1.-,2.1.1.43); Methyl-TRANSFERASE INHIBITOR complex, TRANSFERASE-TRANSFERASE INHIBITOR; HET: SAM, 7L6; 2.6A {Homo sapiens} SCOP: b.85.7.0; Related PDB entries: 5TUY_A | 98.13 | 0.0000016 | 3.3 | 115 | 268 |
| 3F9X_B | Histone-lysine N-methyltransferase SETD8 (E.C.2.1.1.43), Histone; methyltransferase, histone, SET, lysine, Alternative; HET: MLY, SAH; 1.25A {Homo sapiens}; Related PDB entries: 1ZKK_B 1ZKK_A 1ZKK_D 1ZKK_C 3F9W_B 3F9W_A 3F9W_D 3F9W_C 3F9X_A 3F9X_D 3F9X_C 3F9Y_B 3F9Y_A 5W1Y_B | 98.12 | 0.0000018 | 2.9 | 101 | 166 |
| 5VSC_B | Histone-lysine N-methyltransferase EHMT2 (E.C.2.1.1.-,2.1.1.43); protein-small molecule inhibitor complex, TRANSFERASE-TRANSFERASE; HET: SAM, 9HJ; 1.4A {Homo sapiens} SCOP: b.85.7.0; Related PDB entries: 3HNA_B 3HNA_A 5TTG_B 5TTG_A 5V9J_B 5V9J_A 4I51_B 4I51_A 3SW9_B | 98.11 | 0.0000011 | 1.7 | 105 | 275 |

#### Zn finger

| | | | | | | |
|---|---|---|---|---|---|---|
| 1VD4_A | Transcription initiation factor IIE, alpha; ZINC FINGER, Transcription; HET: ZN; NMR {Homo sapiens} SCOP: g.41.3.1 | 87.21 | 0.43 | 1.9 | 27 | 62 |
| 4AYB_P | DNA-DIRECTED RNA POLYMERASE (E.C.2.7.7.6); TRANSFERASE, MULTI-SUBUNIT, TRANSCRIPTION; HET: ZN; 3.202A {SULFOLOBUS SHIBATAE}; Related PDB entries: 3HKZ_X 2Y0S_P 4V8S_BP 2WB1_X 4V8S_AX 2WAQ_P 3HKZ_P 2PMZ_P 2Y0S_X 2PMZ_Z 2WB1_P | 86.97 | 0.29 | 0.7 | 31 | 48 |
| 4QIW_W | DNA-directed RNA polymerase (E.C.2.7.7.6), DNA-directed; Transcription, DNA-directed RNA polymerase; HET: ZN; 3.5A {Thermococcus kodakarensis}; Related PDB entries: 4QIW_P | 86.25 | 0.6 | 2 | 34 | 49 |
| 5K2M_K | RimK-related lysine biosynthesis protein, Probable; ATP-dependent amine/thiol ligase family Amino-group; HET: SO4, ADP, PO4, UN1; 2.18A {Thermococcus kodakarensis (strain ATCC BAA-918 / JCM 12380 / KOD1)}; Related PDB entries: 5K2M_F 5K2M_N 5K2M_E 5K2M_L 5K2M_M | 82.62 | 0.95 | 1.7 | 47 | 53 |

### Figure S3E

#### Chromo-J3C

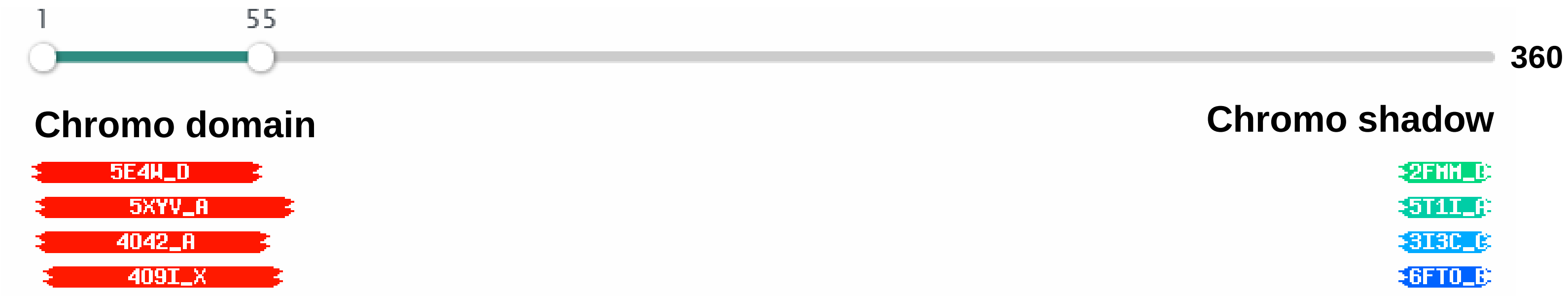


#### Chromo domain

| Hit | Name | Probability | E-value | SS | Cols | Target Length |
|---|---|---|---|---|---|---|
| 5E4W_D | Thioredoxin-1, Signal recognition particle 43; Signal recognition particle, chromodomain, membrane; HET: GOL; 2.8A {Escherichia coli O157:H7}; Related PDB entries: 5E4W_C | 98.52 | 1.1e-7 | 4.2 | 49 | 105 |
| 5XYV_A | RHINO, Protein deadlock; piRNA pathway, Chromoshadow, Complex, PROTEIN; 2.1A {Drosophila melanogaster}; Related PDB entries: 5XYV_B | 98.07 | 0.0000015 | 1.4 | 58 | 78 |
| 4O42_A | Chromodomain-helicase-DNA-binding protein 1 (E.C.3.6.4.12), Nonstructural; viral, chromodomain, Structural Genomics, Structural; HET: MLY, UNX; 1.87A {Homo sapiens}; Related PDB entries: 4NW2_A 4NW2_C 2B2V_B 2B2V_A 2B2W_B 2B2W_A 2B2T_B 2B2T_A 2B2U_B 2B2Y_B 2B2U_A 2B2Y_A | 97.99 | 0.0000054 | 3.6 | 54 | 194 |
| 4O9I_X | Chromodomain-helicase-DNA-binding protein 4 (E.C.3.6.4.12); CHD4 double chromodomains, HYDROLASE; HET: MSE; 2.6A {Homo sapiens} | 97.84 | 0.0000097 | 2.6 | 57 | 202 |

#### Chromo shadow

| | | | | | | |
|---|---|---|---|---|---|---|
| 2FMM_D | Protein EMSY, Chromobox protein homolog; ENT DOMAIN, CHROMO SHADOW DOMAIN; HET: SO4; 1.8A {Homo sapiens} SCOP: b.34.13.2; Related PDB entries: 1S4Z_B 1S4Z_A 2FMM_B 2FMM_A 2FMM_C 1DZ1_A 1DZ1_B | 92.35 | 0.066 | 1 | 52 | 74 |
| 5T1I_A | Chromobox protein homolog 3, histone-H3; Structural Genomics, Structural Genomics Consortium; HET: UNX, N7P, AAR; 1.6A {Homo sapiens}; Related PDB entries: 5T1I_B 6HW2_C 3KUP_B 3KUP_D 3KUP_A 3KUP_C | 60.15 | 7.8 | 2.1 | 21 | 68 |
| 3I3C_C | Chromobox protein homolog 5; CBX5, Chromo Shadow Domain, Structural; 2.48A {Homo sapiens} SCOP: b.34.13.2; Related PDB entries: 3I3C_B 3I3C_D 3I3C_A | 54.34 | 12 | 2.3 | 21 | 75 |
| 6FTO_B | Chromo domain-containing protein 2, Chromatin; chromoshadow domain, complex, chromatin remodeler; HET: HEZ; 1.6A {Schizosaccharomyces pombe}; Related PDB entries: 6FTO_A | 48.38 | 19 | 2.5 | 21 | 66 |

### Figure S3F

## CS-J3C

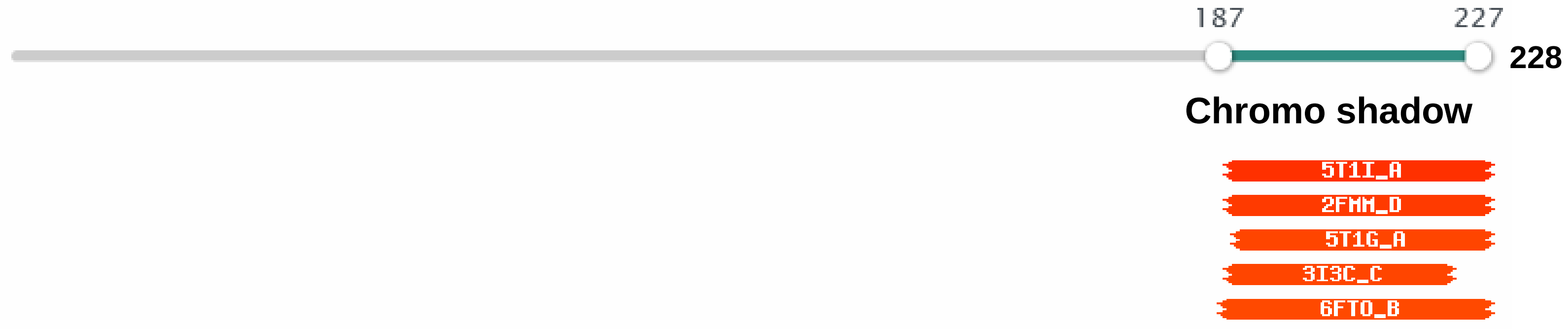


#### Chromo shadow

| Hit | Name | Probability | E-value | SS | Cols | Target Length |
|---|---|---|---|---|---|---|
| 5T1I_A | Chromobox protein homolog 3, histone-H3; Structural Genomics, Structural Genomics Consortium; HET: UNX, N7P, AAR; 1.6A {Homo sapiens}; Related PDB entries: 5T1I_B 6HW2_C 3KUP_B 3KUP_D 3KUP_A 3KUP_C | 95.88 | 0.011 | 3.3 | 41 | 68 |
| 2FMM_D | Protein EMSY, Chromobox protein homolog; ENT DOMAIN, CHROMO SHADOW DOMAIN; HET: SO4; 1.8A {Homo sapiens} SCOP: b.34.13.2; Related PDB entries: 1S4Z_B 1S4Z_A 2FMM_B 2FMM_A 2FMM_C 1DZ1_A 1DZ1_B | 95.12 | 0.027 | 3.2 | 41 | 74 |
| 5T1G_A | Chromobox protein homolog 1, H31; chromo shadow domain, structural genomics; 1.9A {Homo sapiens}; Related PDB entries: 3Q6S_B 3Q6S_A 3Q6S_D 3Q6S_C 6HW2_B | 94.18 | 0.061 | 3.1 | 40 | 79 |
| 3I3C_C | Chromobox protein homolog 5; CBX5, Chromo Shadow Domain, Structural; 2.48A {Homo sapiens} SCOP: b.34.13.2; Related PDB entries: 3I3C_B 3I3C_D 3I3C_A | 94.16 | 0.059 | 3.1 | 35 | 75 |
| 6FTO_B | Chromo domain-containing protein 2, Chromatin; chromoshadow domain, complex, chromatin remodeler; HET: HEZ; 1.6A {Schizosaccharomyces pombe}; Related PDB entries: 6FTO_A | 93.78 | 0.086 | 3.2 | 42 | 66 |

#### Figure S3G

##### HPC-J3C (Tb927.4250)

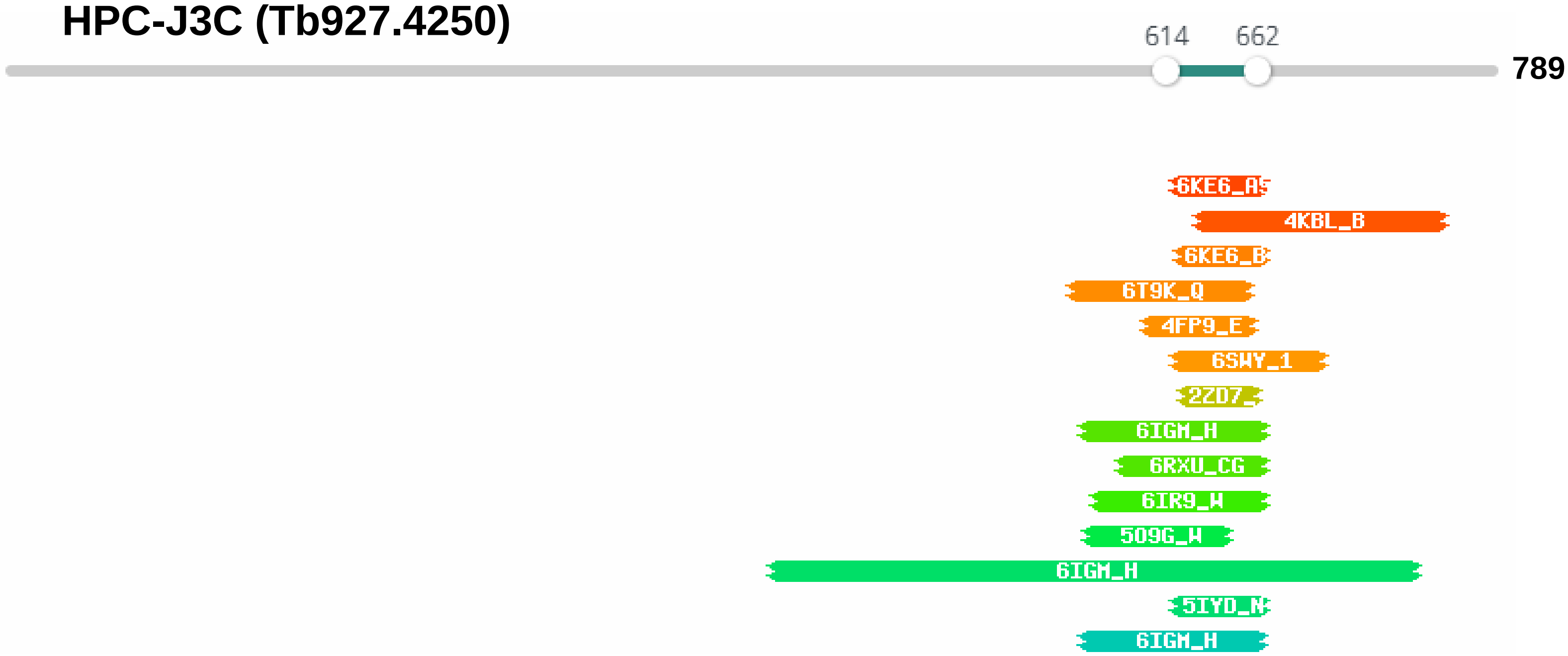


| Nr | Hit | Name | Probability | E-value | Score | SS | Aligned cols | Target Length |
|---|---|---|---|---|---|---|---|---|
| 1 | 6KE6_A5 | 40S ribosomal protein S1-A, 40S; ribosome assembly, 90S pre-ribosome, cryo-EM; HET: GTP; 3.4A {Saccharomyces cerevisiae | 94.15 | 0.047 | 55.57 | 3.4 | 49 | 643 |
| 2 | 4KBL_B | E3 ubiquitin-protein ligase ARIH1 (E.C.6.3.2.-); RING-IBR-RING, E3 ubiquitin ligase, LIGASE; HET: ZN; 3.3A {Homo sapiens | 92.76 | 0.22 | 50.65 | 5.6 | 91 | 559 |
| 3 | 6KE6_B1 | 40S ribosomal protein S1-A, 40S; ribosome assembly, 90S pre-ribosome, cryo-EM; HET: GTP; 3.4A {Saccharomyces cerevisiae | 88.99 | 0.36 | 50.97 | 3 | 47 | 923 |
| 4 | 6T9K_Q | Transcription factor SPT20, Protein SPT3; Coactivator, Transcription, Histone acetyltransferase, Histone; 3.3A {Saccharo | 88.17 | 1.2 | 50.31 | 6.5 | 76 | 657 |
| 5 | 4FP9_E | Putative methyltransferase NSUN4 (E.C.2.1.1.-), mTERF; Modification enzyme, TRANSFERASE; HET: SAM, SO4; 2.9A {Homo sapie | 87.83 | 0.88 | 42.65 | 4.4 | 59 | 335 |
| 6 | 6SWY_1 | Vacuolar import and degradation protein; Suppressed, Suppreseed, LIGASE; 3.2A { Saccharomyces cerevisiae YJM1133} | 87.23 | 0.92 | 52.23 | 5 | 79 | 1064 |
| 7 | 2ZD7_A | Vacuolar protein sorting-associated protein 75; Histone chaperone, vps75, NAP1, Nucleus; 1.85A {Saccharomyces cerevisiae | 81.27 | 1.8 | 40.93 | 3.4 | 41 | 264 |
| 8 | 6IGM_H | RuvB-like 1 (E.C.3.6.4.12), RuvB-like 2; SRCAP complex, TRANSCRIPTION; 4.0A {Homo sapiens} | 75.06 | 2.7 | 53.85 | 3.3 | 97 | 3230 |

### Figure S3H

#### LEO1

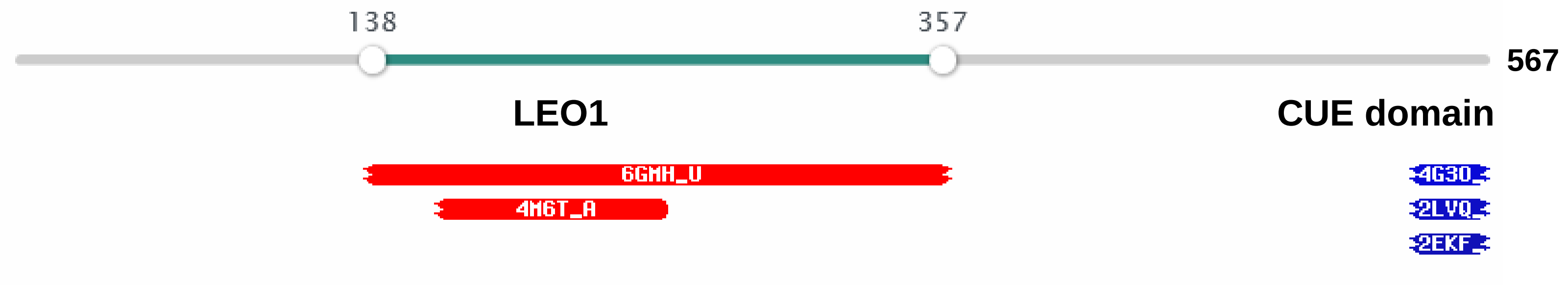


#### LEO1

| Hit | Name | Probability | E-value | SS | Cols | Target Length |
|---|---|---|---|---|---|---|
| 6GMH_U | RPB1, DNA-directed RNA polymerase subunit; DNA, RNA Polymerase, DSIF, PAF1c; HET: SEP, TPO; 3.1A {Sus scrofa} | 99.97 | 8.1e-31 | 13.2 | 188 | 776 |
| 4M6T_A | RNA polymerase II-associated factor 1; Paf1-Leo1 subcomplex, transcription elongator, Transcription; HET: SAM; 2.498A {Homo sapiens} | 99.85 | 2e-21 | 8.3 | 81 | 183 |

#### CUE

| | | | | | | |
|---|---|---|---|---|---|---|
| 4G3O_A | --REMARK 3; all-helical structure, ligase, BAG6; 1.6A {Homo sapiens}; Related PDB entries: 2EJS_A | 37.21 | 68 | 4.2 | 28 | 58 |
| 2LVQ_D | Ubiquitin, E3 ubiquitin-protein ligase AMFR; CUE domain, SIGNALING PROTEIN-LIGASE complex; NMR {Homo sapiens}; Related PDB entries: 2LVN_C 2LVO_C 2LVP_C | 35.75 | 77 | 4.2 | 28 | 52 |
| 2EKF_A | Ancient ubiquitous protein 1; CUE, Ubiquitin ligase complex, Ubiquitin-conjugating; NMR {Homo sapiens} | 34.55 | 80 | 4.3 | 28 | 61 |

### Figure S3I

#### CDC73

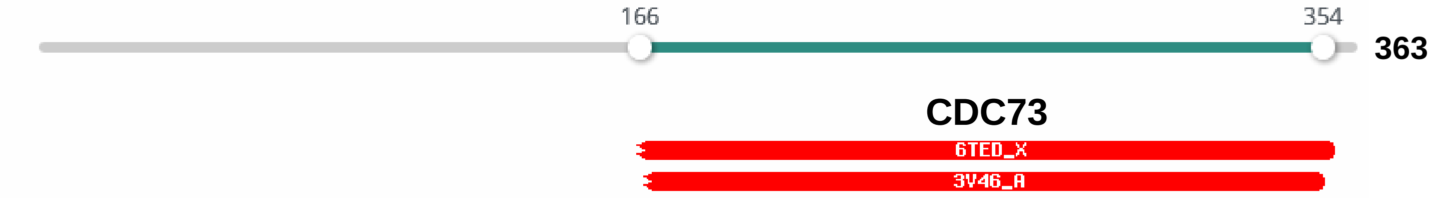


#### CDC73

| Hit | Name | Probability | E-value | SS | Cols | Target Length |
|---|---|---|---|---|---|---|
| 6TED_X | DNA-directed RNA polymerase subunit, DNA-directed; Polymerase, elongation complex, RNA, DNA; HET: SEP, TPO;{Sus scrofa} | 100 | 2.2e-43 | 17.7 | 175 | 531 |
| 3V46_A | Cell division control protein 73; Ras-like fold, non-GTP binding, Protein; 1.549A {Saccharomyces cerevisiae}; Related PDB entries: 4DM4_A 4DM4_B | 100 | 3.4e-43 | 16.4 | 160 | 170 |

### Figure S3J

#### CTR9

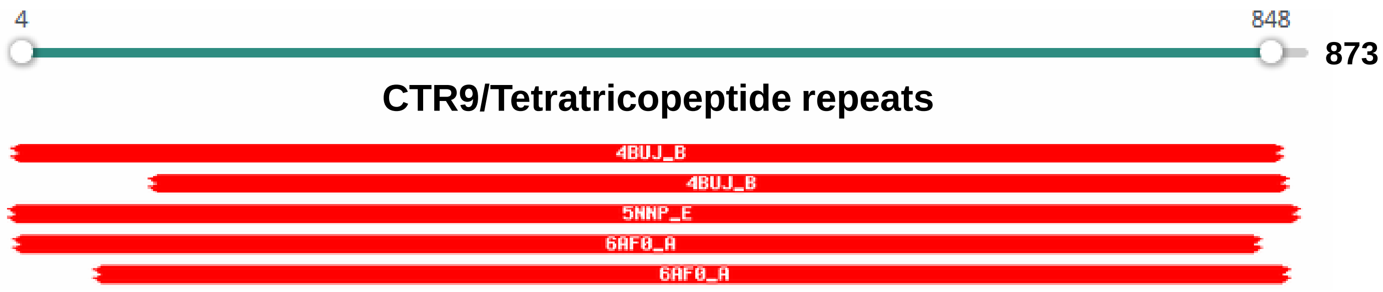


#### CTR9/Tetratricopeptide repeats

| Hit | Name | Probability | E-value | SS | Cols | Target Length |
|---|---|---|---|---|---|---|
| 4BUJ_B | ANTIVIRAL HELICASE SKI2 (E.C.3.6.4.13), SUPERKILLER; HYDROLASE, DEXH BOX HELICASE, RNA; HET: SO4; 3.7A {SACCHAROMYCES CEREVISIAE}; Related PDB entries: 4BUJ_F 5MC6_i | 100 | 1.9e-44 | 89.5 | 784 | 1436 |
| 4BUJ_B | ANTIVIRAL HELICASE SKI2 (E.C.3.6.4.13), SUPERKILLER; HYDROLASE, DEXH BOX HELICASE, RNA; HET: SO4; 3.7A {SACCHAROMYCES CEREVISIAE}; Related PDB entries: 4BUJ_F 5MC6_i | 100 | 1.7e-41 | 77.7 | 695 | 1436 |
| 5NNP_E | N-terminal acetyltransferase-like protein, Naa10, HypK; N-acetylation, NATs, Naa15, Naa10, HypK; HET: CMC, PO4, FME, GOL; 2.602A {Chaetomium thermophilum (strain DSM 1495 / CBS 144.50 / IMI 039719)}; Related PDB entries: 5NNR_A 5NNR_D 5NNP_A | 100 | 3.1e-41 | 66.3 | 693 | 745 |
| 6AF0_A | Ctr9 protein, Paf1 protein, Cdc73; Transcription elongation Paf1 ; 2.88A {Myceliophthora thermophila (strain ATCC 42464 / BCRC 31852 / DSM 1799)} | 100 | 8.4e-39 | 85.2 | 735 | 939 |
| 6AF0_A | Ctr9 protein, Paf1 protein, Cdc73; Transcription elongation Paf1 ; 2.88A {Myceliophthora thermophila (strain ATCC 42464 / BCRC 31852 / DSM 1799)} | 100 | 9.8e-38 | 76.1 | 675 | 939 |

### Figure S3K

#### DCNL

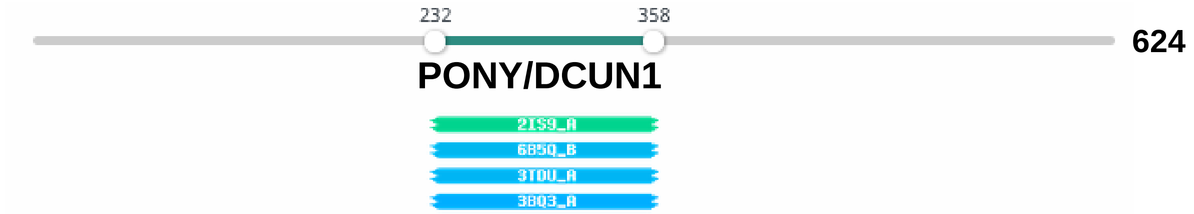


| Hit | Name | Probability | E-value | SS | Cols | Target Length |
|---|---|---|---|---|---|---|
| 2IS9_A | Defective in cullin neddylation protein; ubiquitin, DCN1, TRANSCRIPTION; HET: MES; 1.92A {Saccharomyces cerevisiae}; Related PDB entries: 3O6B_I 3O6B_A 3O6B_C 3O6B_E 3O6B_G 3O2P_A 3TDI_B 3TDI_A | 61.6 | 64 | 9.1 | 102 | 204 |
| 6B5Q_B | DCN1-like protein 1; E3 ligase, complex, Ligase-Inhibitor complex; HET: MLY, PPI, CZS, 2KY, PGE, 1XY; 2.16A {Homo sapiens}; Related PDB entries: 5UFI_B 5UFI_D 6B5Q_A 5UFI_A 5UFI_C | 55.91 | 43 | 7 | 105 | 225 |
| 3TDU_A | DCN1-like protein 1, Cullin-1, NEDD8-conjugating; E2:E3, Ligase-protein binding complex; 1.5A {Homo sapiens}; Related PDB entries: 4GAO_B 4GAO_A 4GAO_D 4GAO_G 4P5O_F 3TDZ_B 3TDZ_A 4P5O_E 3TDU_B | 55.56 | 39 | 6.6 | 105 | 200 |
| 3BQ3_A | Defective in cullin neddylation protein; ubiquitin, Nedd8, neddylation, ubiquitination, SCF; HET: MSE; 1.9A {Saccharomyces cerevisiae} | 54.65 | 71 | 8.4 | 102 | 270 |

### Figure S3L

#### RTFL

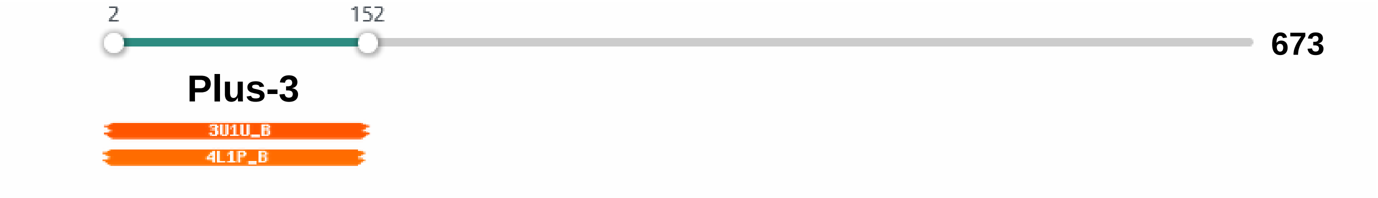


| Hit | Name | Probability | E-value | SS | Cols | Target Length |
|---|---|---|---|---|---|---|
| 3U1U_B | RNA polymerase-associated protein RTF1 homolog; PLUS-3, transcription elongation, Structural Genomics; HET: SO4, MSE, UNX; 1.8A {Homo sapiens} SCOP: b.34.21.1; Related PDB entries: 3U1U_A | 92.84 | 0.048 | 1 | 109 | 137 |
| 4L1P_B | RNA polymerase-associated protein RTF1 homolog; Tutor, Plus3, Peptide binding protein; HET: GOL; 2.12A {Homo sapiens} SCOP: l.1.1.1, b.34.21.1; Related PDB entries: 2BZE_A 2DB9_A 4L1U_B 4L1U_F 4L1U_A 4L1U_D 4L1U_C 4L1U_E 4L1P_A | 90.86 | 0.095 | 0.6 | 108 | 138 |
